# Dissecting Agronomically Favorable Genotypes in Temperate Japonica Rice via Haplotype Analysis of a Japan-MAGIC Population

**DOI:** 10.64898/2025.12.04.692267

**Authors:** Hirofumi Fukuda, Akari Fukuda, Toshihiro Sakamoto, Yoshihiro Kawahara, Ken Naito, Taiji Kawakatsu, Jun-ichi Yonemaru, Daisuke Ogawa

## Abstract

- Crop breeding assembles genomic variants into cultivars via crossing and selection. Phenotypic selection has improved yield and lodging tolerance but has limited genetic insights. We show how specific genomic variants and their combinations underpin advances in modern rice breeding in Japan.
- Through genome-wide association study using a multi-parent advanced-generation intercross population derived from four temperate *japonica* cultivars, we identified 11 quantitative trait loci (QTLs) for key agronomic traits, including days to heading, shoot biomass, panicle length, and culm length under field conditions. *GA20ox1, GA20ox2*, and *Hd1* were among the QTLs, and their natural variants were well conserved in temperate *japonica* cultivars bred in Japan, underscoring distinct selection pressures at these loci.
- By integrating genotype data with 5-year yield-performance-evaluation trials of elite cultivars, we found that cultivars carrying multiple-copy *GA20ox1* and functional *Hd1*, together with non-functional *ga20ox2*, tended to have shorter culms and higher grain yield than cultivars with multiple-copy *GA20ox1*, functional *Hd1*, and *GA20ox2*. This yield advantage was consistent across latitudes in Japan.
- These results reveal favorable genotype combinations underlying modern *japonica* improvement and provide a genomic framework for breeding semi-dwarf, high-yielding cultivars adapted to temperate rice-growing regions in Asia.

## Introduction

Rice (*Oryza sativa* L.) is a staple food for more than half of the global population, particularly in temperate regions, including Japan, making its genetic improvement a global priority for ensuring food security under increasing environmental and demographic pressures. To meet rising food demand, breeding programs have developed cultivars that not only have high yield potential but are also resistant to lodging owing to their semi-dwarf phenotypes (Hargrove & Cabanilla 1979; Hargrove *et al*., 1988). Landmark cultivars such as IR8, developed by International Rice Research Institute in the 1960s, and the Japanese cultivar Reimei, derived from gamma-ray-induced mutants of Fujiminori, played pivotal roles in enhancing lodging resistance through semi-dwarf phenotypes and served as key parental lines in subsequent Japanese breeding efforts (Futsuhara *et al*., 1967; Ashikari *et al*., 2002). The semi-dwarf characteristics of these cultivars were later attributed to distinct mutations in *Sd1*/*GA20ox2*, which encodes a key enzyme in the late stage of gibberellin biosynthesis (Ashikari *et al*., 2002; Spielmeyer *et al*., 2002). However, rice breeding in Japan from the 1960s to early 2000s relied heavily on empirical phenotypic selection, limiting insights into the genetic basis of agronomic traits. To advance breeding strategies and improve elite cultivars, it is essential to identify the genetic variants that govern key agronomic traits, including semi-dwarf architecture.

Dwarfism in rice can depress canopy photosynthesis and decrease biomass, because total leaf area is reduced and leaves are packed more densely, thus increasing self-shading and lowering light-use efficiency (reviewed by Weng *et al*., 2025). As canopy photosynthesis reflects the integrated performance of all leaves, leaf size and spatial arrangement determine how much leaf area can be deployed and how effectively light is intercepted. Semi-dwarf plants carrying the *ga20ox2* allele typically have shorter leaves and reduced stature (Sasaki *et al*., 2002; Feng *et al*., 2023), and this can constrain photosynthetic capacity in dense canopies. Breeding practice therefore likely favored *ga20ox2*-possessing lines that maintain leaf length and canopy height (CH) after crossing with core semi-dwarf cultivars. In Japan, culm length (CL) has remained broadly constant for decades, whereas biomass and grain yield have increased (Matsushita *et al*., 2024). This trend indicates that breeders consequently paired *ga20ox2* alleles with genotypes that sustain shoot growth, on the basis of phenotypic selection, thus achieving higher biomass and yield without sacrificing lodging resistance.

Genome-wide association studies (GWAS) using rice populations have been instrumental in identifying quantitative trait loci (QTLs) associated with flowering time, plant architecture, yield-related traits, and responses to abiotic and biotic stresses (Yano *et al*., 2016; Sertse *et al*., 2021; Serrie *et al*., 2025). These studies highlight the power of high-resolution mapping to uncover natural variations relevant to breeding. However, the use of highly structured natural accessions can introduce confounding due to population stratification (Lander & Schork, 1994; Cardon & Palmer, 2003) and cryptic relatedness (Sillanpää *et al*., 2011), potentially leading to spurious associations. To address these limitations, multi-parental populations such as multi-parent advanced-generation intercrosses (MAGIC) have emerged as powerful resources for the genetic dissection of complex traits (McMullen *et al*., 2009; reviewed by Huang *et al*., 2015; Abdelraheem *et al*., 2021). MAGIC populations are generated through controlled intercrossing among multiple founder lines, thus helping to minimize hidden population structure and maximize recombination. In this way, they combine broad genetic diversity with high mapping resolution and reduced confounding effects. Previously, we conducted haplotype-based GWAS by using a mixed *indica*–*japonica* MAGIC population consisting of eight founder lines derived from Japanese cultivars (i.e., Japan-MAGIC1, JAM1), and we successfully identified well-known genes and novel QTLs associated with glutinous endosperm, plant spreading habit, CL, aboveground dry weight (AGW, i.e., shoot biomass), panicle weight (PW), and grain shape (Ogawa *et al*., 2018a, 2018b, 2021). We recently developed a new MAGIC population, Japan-MAGIC2 (JAM2), derived from four elite temperate *japonica* founders bred in Japan in the 21st century (Taniguchi *et al*., 2025a). Two-year field evaluations demonstrated that the JAM2 population consistently had broad phenotypic diversity, indicating that it harbors a wide range of natural variants underlying modern elite temperate *japonica* cultivars.

In this study, we aimed to identify which natural variants have been used throughout the Japanese breeding history of temperate *japonica* cultivars and to elucidate their roles in the development of modern Japanese cultivars. To explore the natural variants, we first conducted haplotype-based GWAS by using the JAM2 population and identified haplotypes/QTLs associated with agronomic traits important for lodging resistance and yield. We then conducted data mining of DNA sequencing results, including whole-genome sequencing, to identify the natural variations in candidate genes underlying the identified QTLs. Finally, by using Japanese historical field records of elite temperate *japonica* cultivars, we validated the roles of combinational genotypes of these candidates in agronomic traits important for lodging resistance and yield.

## Materials and Methods

### Development and cultivation of JAM2 lines

JAM2 lines were developed as described previously (Taniguchi *et al*., 2025a). Briefly, four Japanese cultivars— Akidawara (AD), Iwaidawara (ID), Tachiharuka (TH), and Toyomeki (TM)—were used as the parents of JAM2 on the basis of their genetic diversity and regional breeding histories across Japan (Fig. 1a, Fig. S1). We first crossed AD with TM and ID with TH to produce seeds of two types. Then these hybrids were crossed to produce four-way recombinants. By using the single-seed descent method, JAM2 were developed. One hundred JAM2 lines (F5 and F6) in 2022 and 2023 were used in this study. Seeds soaked in water at 30 °C for 2 days were sown in trays filled with soil and incubated at 30 °C in the dark for 2 days. Seedlings were grown in a paddy field in Kannondai, Tsukuba, for a month, and then 33 seedlings per line were transplanted (11 plants 18 cm apart × 3 rows 30 cm apart, no replicates) into a nearby paddy field and cultivated for 5 months. The date of sowing was 18 April in 2022 and 17 April in 2023.

**Fig. 1.**
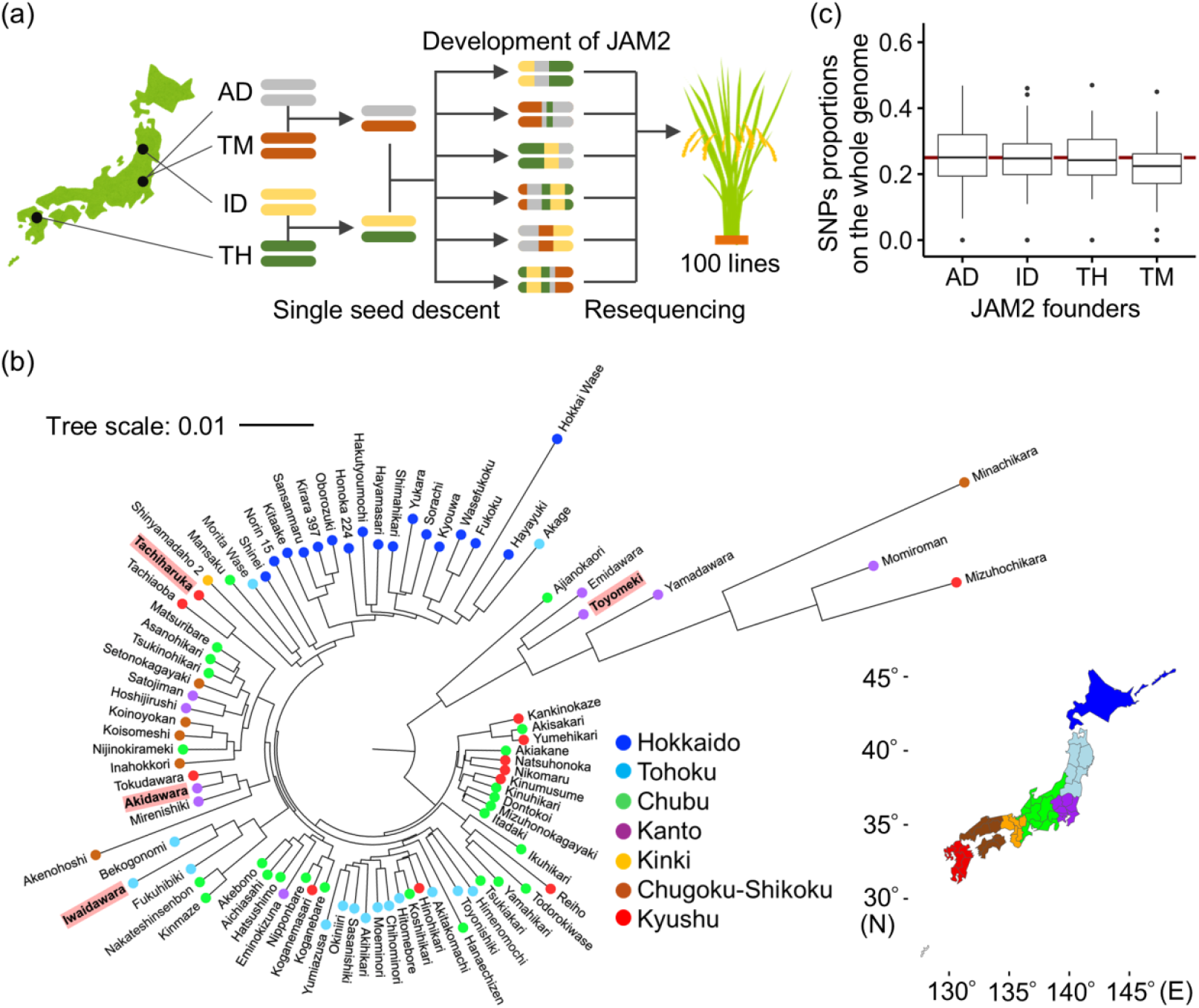
Genomic information on JAPAN-MAGIC2 founders and the JAPAN-MAGIC2 population. (a) Schematic overview of the process used to establish the JAM2 population, consisting of 100 lines derived from four temperate *japonica* cultivars: Akidawara (AD), Iwaidawara (ID), Tachiharuka (TH), and Toyomeki (TM). (b) Phylogenetic tree of temperate *japonica* rice cultivars bred in Japan. Colored circles indicate the regions where each cultivar was developed. The four founder cultivars of the JAM2 population are highlighted. (c) Proportions of JAM2-founder-derived single nucleotide polymorphisms (SNPs) in the whole genome of JAM2 lines; 131,537 SNPs were used for this analysis.

### Haplotype analysis

Haplotypes among the 100 lines of the JAM2 population were estimated by using a haplotype-based approach similar to that used in Arabidopsis MAGIC populations (Kover *et al*., 2009). The estimation was further refined on the basis of methodologies developed for the Japan-MAGIC rice population (Ogawa *et al*., 2018a, 2018b, 2021). The whole genome was resequenced by using short-read sequencing technology. Single nucleotide polymorphisms (SNPs) were detected and filtered by using the bioinformatics pipelines described by Ogawa *et al*. (2018a, 2018b, 2021). Haplotypes at each SNP locus were defined on the basis of the genotypes of the four founders. Haplotype blocks were inferred by grouping contiguous SNPs that had consistent founder-origin patterns.

### Phenotypic measurement and quantification

Days to heading (DTH) was scored as the number of days from transplanting rice to the field to the appearance of the first panicle in more than half of the plants in each JAM2 line. CL and panicle length (PL) of the longest culm on each plant were measured with a ruler, and panicle number (PN) was counted, from 10 days to a month after heading. To determine PW and stem and leaf weight (SLW), shoots of mature plants were air-dried for 1 to 2 months in a drying room and then cut 3 cm below the panicle base to separate the parts; the AGW was calculated as PW + SLW. The averages of five plants in the middle lane, except for the plants at both edges, were used as the CL, PW, SLW, and AGW values of each JAM2 line. CH was quantified by using images captured by an unmanned aerial vehicle (UAV) in our previous study (Taniguchi *et al*., 2025a) and used in this study. We analyzed historical field records collected by NARO for yield performance evaluation from 2017 to 2021 for 52 Japanese cultivars evaluated at up to 69 field stations across the Japanese archipelago, including part of the data used in previous studies (Matsushita *et al*., 2024; Taniguchi *et al*., 2025b). CL, shoot biomass, and estimated grain yield (t/ha) were aggregated as means computed within “cultivar × prefecture × year” combinations and used for subsequent analyses.

### RNA extraction

One most expanded leaf per JAM2 line was collected at 6 weeks after transplanting (1 July 2022 and 29 June 2023) from 9:00 to 11:00 am, immediately frozen in liquid N_2_, and stored at −80 °C. Frozen samples were crushed by using a multi-specimen cell-disruption device (BMS-A20TP, Bio Medical Science, Shinjuku, Japan), and total RNA from each sample was extracted immediately by using an RNeasy Plant Mini Kit (Qiagen, Inc., Venlo, the Netherlands) in accordance with the manufacturer’s protocol.

### Library preparation and data processing for transcriptome analysis

RNA-sequencing (RNA-seq) libraries were prepared by using a NEBNext Ultra II Directional mRNA-seq kit for Illumina (New England Biolabs, Inc., Ipswich, USA) in accordance with the manufacturer’s protocols. The libraries were sequenced by using Illumina NovaSeq X Plus on a 25B flow cell with paired-end 150-bp and unique dual-index reads. The reads were preprocessed in CLC Genomics Workbench version 25.0 (Qiagen). Reads were first trimmed with the following parameters: quality limit, 0.05; maximum number of ambiguities, 2. Next, raw reads were mapped to the genome assembly and gene sets of “Os-Nipponbare-Reference-IRGSP-1.0” with the following parameters: mismatch cost, 2; insertion–deletion cost, 3; length fraction, 0.8; similarity fraction, 0.8; and maximum number of hits per read, 10. Then, uniquely mapped reads were counted. Genes with at least one raw read count in all of the library samples for each comparison were analyzed. Log_2_(fold change) and adjusted *P-*values (Wald test using the “DESeq” function) were calculated from raw read counts by using the R package DESeq2 (version 1.42.1) in R software (version 4.3.2). Differentially expressed genes compared with the control group were extracted on the basis of adjusted *P-* values (padj) < 0.05.

### Haplotype-based GWAS and expression GWAS

Haplotype-based GWAS used the haplotype information at each SNP and phenotype data in non-parametric ANOVA (Kruskal–Wallis rank sum test) in the “kruskal.test” package of R software. *P*-values obtained from the statistical analysis were used for the Manhattan plot. To identify QTLs for phenotypes, we selected SNPs with *P-*values < 1.0 × 10^−3^. In the case of consecutive selected SNPs, the SNP with the lowest *P*-value defined the QTL unless the distance between them was more than 2 Mb. For the expression GWAS, the haplotype information and transcriptome data of 100 JAM2 lines were used. *P*-values for each transcript were obtained from the statistical analysis in the same way as for the GWAS and used for the Manhattan plot for target genes.

### Genomic DNA extraction, size selection, whole-genome sequencing, and assembly to detect tandem repeat structures

Genomic DNA was extracted from each JAM2 founder by using a Nucleobond HMW DNA Kit (Takara Bio., Inc., Kusatsu, Japan) in accordance with the manufacturer’s instructions. To enrich the extracts for high–molecular weight DNA and reduce the number of short fragments, the extracts were subjected to size selection with a Short Read Eliminator Kit (Pacific Biosciences, Menlo Park, USA), which removes DNA fragments shorter than 10 kb to improve long-read recovery efficiency. For sequencing, 2.5 μg of size-selected genomic DNA was used to construct libraries with a Ligation Sequencing Kit LSK114 (Oxford Nanopore Technologies KK, Tokyo, Japan). The resulting libraries were sequenced on an R10.1 flow cell on PromethION 24 (Oxford Nanopore Technologies KK). The raw POD5 data were transformed into fastq with the dna_r10.4.1_e8.2_400bps_fast@v5.0.0 model in Dorado 0.9.0 (https://github.com/nanoporetech/dorado). To obtain better quality reads, we used the default parameters for cultivar AD, and we used the SeqKit2 (Shen *et al*., 2024) with the command “seqkit seq -m 3000 -Q 10” to filter a minimum length of 3000 bases, with an average quality score of at least 10 for cultivars ID, TH, TM. PECAT (https://github.com/lemene/PECAT) with the default parameters was used to obtain a draft genome assembly. The contigs were polished with the SeqKit tool with the command “-m 1000000” to filter a minimum length of 1,000,000 bp. We used Os-Nipponbare-Reference-IRGSP-1.0 (*O. sativa* L. ssp. *japonica* ‘Nipponbare’) as a reference genome assembly.

### Genotyping based on SNPs, DNA short-read depth, and PCR-based assay

Among the QTLs detected, we determined the genotypes for each candidate gene (described later) related to agronomic traits on the basis of neighboring SNPs (Table S1) that constructed haplotype differences in the JAM2 lines: the SNP at M5483 (Chr01: 5,273,347) for *Gn1a*, M27769 (Chr03: 1,299,595) for *SOC1*, M31650 (Chr03: 36,136,637) for *GA20ox1*, M14158 (Chr01: 38,380,261) for *GA20ox2*, and M53254 (Chr06: 9,333,525) for *Hd1*. To confirm the mutations—insertion, deletion, and repeated genomic structures—we performed a PCR-based assay by using established primers (Table S2) for *Gn1a, GA20ox1, GA20ox2*, and *Hd1*. The allele difference in *SOC1* was confirmed by using the TASUKE+ multiple genome browser in RAP-DB (the Rice Annotation Project Database) (https://rapdb.dna.affrc.go.jp/). Next, we estimated the functional differences of each of the *GA20ox1, GA20ox2*, and *Hd1* genotypes. For *GA20ox1*, the multiple-copy type was defined as a functionally strong genotype, either with a locus-to-whole-genome depth ratio greater than 2 (based on short-read sequencing), or with a repeated genomic structure of the *GA20ox1* locus confirmed by PCR assay. Otherwise, it was considered a single copy. For *GA20ox2*, the non-functional alleles carried a >380-bp (length) deletion in the first exon, which originated from the cultivar Dee-geo-woo-gen (Chandler 1968; Ashikari *et al*., 2002). For *Hd1*, the non-functional alleles were characterized by a 36-bp (length) insertion in the first exon. These deletion- and insertion-type mutations in *GA20ox2* and *Hd1*, respectively, can be recognized either in the TASUKE+ browser by using short-read depth information or by PCR-based assay. Using 65 Japanese cultivars for which the year of name release and breeding region were recorded, we determined the combinational genotypes of *GA20ox1, GA20ox2*, and *Hd1* according to the criteria described above.

## Results

### Genomic information on founders of the Japan-MAGIC2 (JAM2) population and JAM2 lines

The JAM2 founders (i.e., AD, ID, TH, and TM) originated from distinct pedigrees and were bred in regions from northeastern to southwestern Japan (Figs. 1a,b; S1). We analyzed the whole-genome resequencing data of a total of 100 JAM2 lines (F5 in 2022, F6 in 2023) and performed short-read genome assemblies to detect SNPs. We found 131,537 SNPs in all the chromosomes (Table S1). The proportions of haplotypes derived from each founder were, in descending order, AD, ID, TH, and TM, with an almost equal distribution of approximately 25% from each founder on the whole genome (Fig. 1c). The distribution of haplotypes differed among the founders at an individual chromosome level (Fig. S2). This indicated that various genomic variants were introgressed among the JAM2 lines.

### QTLs associated with agronomic traits in the JAM2 population, and their candidate genes

The JAM2 population showed variations in DTH, CL, PL, PN, PW, SLW, and AGW under experimental field conditions in 2022 and 2023 (Fig. S3a), as partly described in our previous study (Taniguchi *et al*., 2025a). Haplotype-based GWAS revealed 11 QTLs [4 > –log_10_(*P*) in both 2022 and 2023] related to DTH, CL, PL, SLW, and AGW (Fig. S3b,c, Table 1). Among the QTLs, some haplotypes included loci of agronomically important genes: *Grain number 1a* (*Gn1a*, LOC_Os01g10110/Os01g0197700) (Kim *et al*., 2016; Li *et al*., 2016; Gouda *et al*., 2020) for PL; *Suppressor of overexpression of constans 1* (*SOC1*, LOC_Os03g03070/Os03g0122600) (Lee *et al*., 2004) for DTH, *Heading date 1* (*Hd1*, LOC_Os06g16370/Os06g0275000) (Yano *et al*., 2000) for DTH, CL, and AGW; and *Gibberellin 20-oxidase 1* (*GA20ox1*, LOC_Os03g63970/Os03g0856700) and *Gibberellin 20-oxidase 2* (*GA20ox2*, LOC_Os01g66100/Os01g0883800) for CL (Table 1). Among these five genes, we found novel natural variants of *Gn1a* (Fig. S4), *SOC1* (Fig. S5), and *GA20ox1* (Fig. 2). We found that newly identified natural variants of *Gn1a* and *SOC1* were used rarely in the past in Japanese breeding of staple rice cultivars (See Supplementary documentation). Variants of *GA20ox1, GA20ox2*, and *Hd1* for CL were used frequently in modern temperate *japonica* cultivars (details described later).

**Table 1.** Identification of putative QTLs for agronomic traits in the JAM2 population.

| (QTL) No | Chr | Position | Trait | 2022 | 2023 | Candidates | Variant reference |
| --- | --- | --- | --- | --- | --- | --- | --- |
| I | 1 | 4958120-4979170 | PL | 1.98E-04 | 6.00E-05 | *Gn1a* | This study |
| II | 1 | 38398436-38513991 | CL | 1.93E-04 | 1.41E-06 | *GA20ox2* | Sasaki et al., 2002 |
| III | 3 | 225264-2403133 | DTH | 1.47E-04 | 1.02E-04 | *SOC1* | This study |
| IV | 3 | 33002789-33034128 | CL | 3.45E-04 | 3.86E-05 |  |  |
| V | 3 | 35099038-35113003 | CL | 1.17E-04 | 2.21E-06 | *GA20ox1* | This study |
| VI | 6 | 7235289-7253747 | DTH | 2.08E-08 | 1.17E-08 |  |  |
| VII | 6 | 9333525-9524731 | DTH | 6.22E-13 | 3.86E-13 | *Hd1* | Yano et al., 2000 |
| VII | 6 | 8987955-9303179 | CL | 7.78E-05 | 4.16E-06 | *Hd1* |  |
| VII | 6 | 9333525-9575842 | AGW | 1.56E-11 | 2.59E-09 | *Hd1* |  |
| VII | 6 | 9315860 | SLW | 5.44E-12 | 6.19E-12 | *Hd1* |  |
| VIII | 6 | 21439006-21452322 | SLW | 1.11E-04 | 4.35E-05 |  |  |
| IX | 6 | 23003215-23056380 | DTH | 1.65E-05 | 1.30E-05 |  |  |
| X | 6 | 23920500 | SLW | 3.95E-05 | 1.63E-05 |  |  |
| XI | 7 | 20826800-20844742 | DTH | 8.60E-04 | 6.70E-04 |  |  |

Summary of putative QTLs detected in this study. Positions indicate the genomic region with $P$ -values $< 0.001$ [ $4 > -\log_{10}(P)$ ] in both 2022 and 2023 among the JAM2 lines. Candidate genes were determined on the basis of their known roles in each agronomic trait in previous studies. DTH, days to heading; CL, culm length; PL, panicle length; PN, panicle number; PW, panicle weight; SLW, stem and leaf dry weight; AGW, aboveground dry weight

**Fig. 2.**
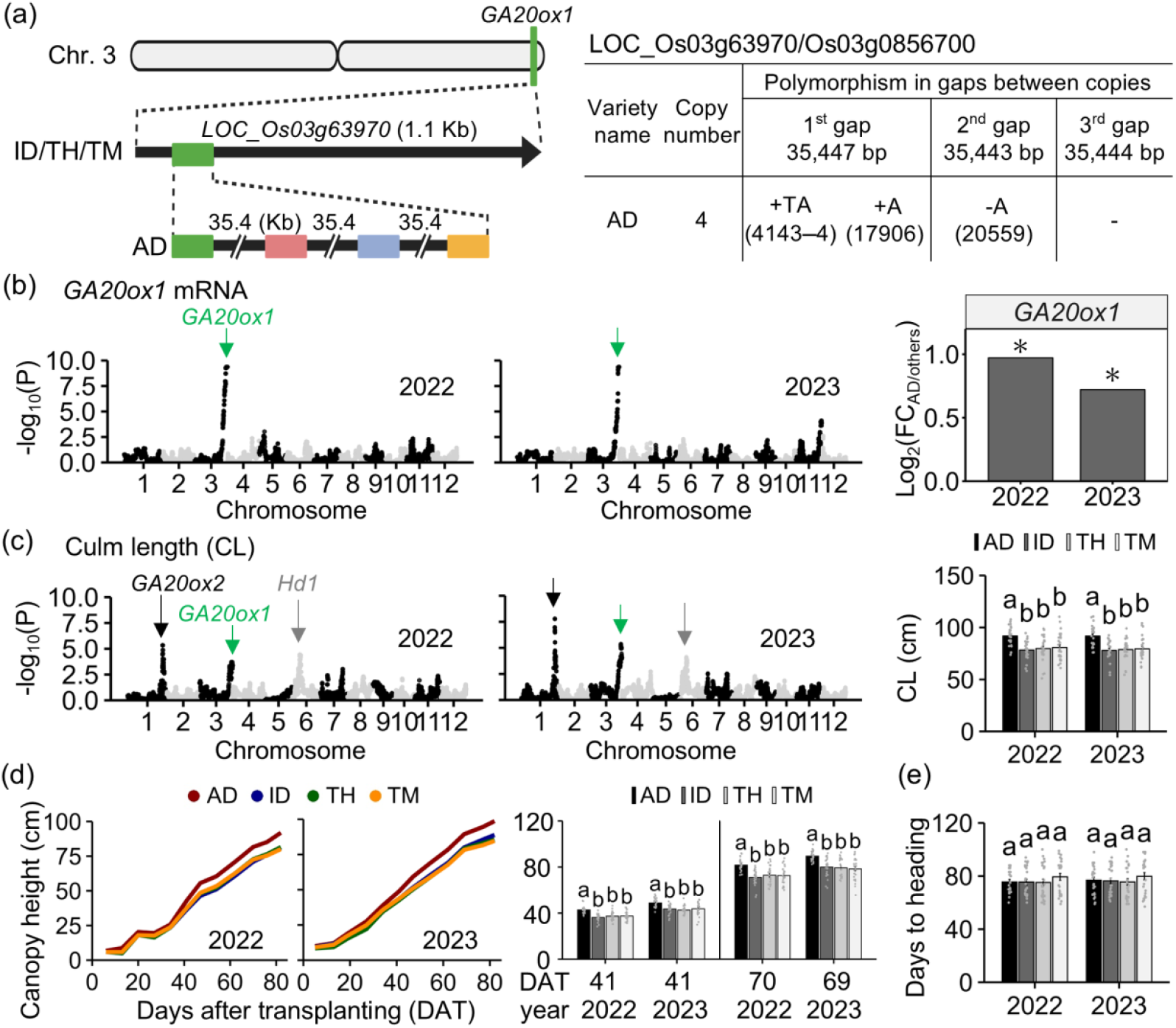
Effect of a putative QTL encoding a new genotype of *GA20ox1* on culm length and canopy height at the vegetative stage. (a) Scheme of the *GA20ox1* (LOC_Os03g63970/Os03g0856700) genotype among the four founders (AD, Akidawara; ID, Iwaidawara; TH, Tachiharuka; TM, Toyomeki). Polymorphisms in three gaps of the *GA20ox1* tandem repeats are described. Numbers in parentheses indicate positions counted from the first nucleotide (+1) in each gap. (b) Expression genome-wide association study (GWAS) of *GA20ox1* among the JAM2 lines. Differences in the expression levels of *GA20ox1* between single and four copies in the JAM2 lines are shown. Asterisks indicate significant differences between genotypes of multiple copies derived from AD and a single copy derived from the other three founders (padj < 0.05, n = 23 to 77). (c) GWAS detected a peak including the *GA20ox1* locus for culm length (CL) among the JAM2 lines. (d) Time series of canopy height (CH) at the vegetative stage and (e) days to heading (DTH) in JAM2 lines classified by the haplotype including the *GA20ox1* locus. Plants were transplanted to experimental fields on 18 May 2022 and 17 May 2023. Time series of CH are shown, with means of each group classified with *GA20ox1* genotypes derived from each JAM2 founder. The means ± SD for CL, CH at two time points, and DTH are shown (n = 23 to 77). Different letters indicate significant differences among JAM2 lines in a haplotype-dependent manner (*P* < 0.05, Tukey–Kramer test). FC, fold change

### Positive role of a natural variant of multiple-copy *GA20ox1* in increasing CH under field conditions

Without SNPs or indels for *GA20ox1* among the JAM2 founders, we conducted long-read sequencing by using high-molecular-weight DNA from each JAM2 founder. We found four tandem copies of *GA20ox1* in AD, whereas the others had only one copy (Fig. 2a). GWAS analysis revealed a single prominent peak near *GA20ox1* (Fig. 2b). When we grouped the JAM2 lines in a haplotype-dependent manner, the mRNA levels of *GA20ox1* in the lines harboring the tandem repeat of *GA20ox1* (*GA20ox1*^AD^) were significantly higher (padj < 0.05) than those in the other lines with a single copy (*GA20ox1*^ID/TH/TM^). These mRNA analyses indicated that the higher copy number of *GA20ox1* was responsible for the high gene expression level among the JAM2 lines. Our GWAS analysis revealed several prominent peaks near *GA20ox1* (Fig. 2c). CL at harvest was significantly longer in *GA20ox1*^*AD*^ than in *GA20ox1*^ID/TH/TM^ among the JAM2 lines. Time-series investigation of CH revealed that the *GA20ox1*^*AD*^ variant significantly increased CH from 41 days after transplanting (DAT) in both 2022 and 2023 (Fig. 2d); this was more than a month before heading, the time of which did not differ among the lines (Fig. 2e). Differences in the *ga20ox2* variant had little effect on CH at 41 DAT in 2022, although the effect of the distinct genotypes was observed at about 70 DAT in both 2022 and 2023, and at 41 DAT in 2023 (Fig. S6a,b). These results suggested that CH differences arose earlier with the multiple-copy *GA20ox1* variant than with the non-functional *ga20ox2* variant.

### Positive effect of *Hd1* on vegetative growth under field conditions

We found a 36-bp insertion in the first exon in TM and a 36-bp insertion with a 43-bp deletion in the first exon in ID (Fig. S7a). These variants had already been reported, and they cause early heading, indicating that the genotype is functionally weak (Yano *et al*., 2000). JAM2 lines harboring haplotypes including the loci of ID- and TM-type *hd1* had significantly shorter DTH, lower AGW, and shorter CL than those of AD- and TH-type *Hd1* (Fig. S7b). Transcriptome analysis showed that the mRNA levels of *early heading date 1* (*Ehd1*, LOC_Os10g32600/Os10g0463400), *heading date 3a* (*Hd3a*, LOC_Os06g06320/Os06g0157700), and *RICE FLOWERING LOCUS T 1* (*RFT1*, LOC_Os06g06300/Os06g0157500) were significantly higher in lines harboring ID- and TM-type *hd1* than in those with AD- and TH-type *Hd1* (Fig. S7c). Together, our results suggested that *Hd1* was a causal gene in the DTH/AGW/CL-related QTLs on chromosome 6 among the JAM2 lines. To validate the exact timing of when the difference in *Hd1* alleles affects shoot biomass, we compared these traits between Koshihikari and a near-isogenic line (NIL) harboring the Kasalath-derived functionally weak *hd1* allele in the Koshihikari genetic background (Itoh *et al*., 2018). Plant height and AGW differences appeared during the period from the heading date of the NIL to that of Koshihikari (Fig. S7d,e). The accumulating evidence indicated that shoot biomass production is enhanced through delayed heading timing controlled by the Hd1–Ehd1–Hd3a/RFT1 pathway.

### Well-conserved genotypes of *GA20ox1, GA20ox2*, and *Hd1* in temperate *japonica* cultivars

Next, we explored the degree of conservation of the natural variants we found in JAM2 (i.e., multiple-copy *GA20ox1*, deletion-type *ga20ox2* allele, insertion-type *hd1* allele) among different rice subspecies. As the long-read sequencing data were quite limited in rice, including Japanese cultivars, in the published dataset (Jain *et al*., 2019; Tanaka *et al*., 2020; Wang *et al*., 2023; Sugimura *et al*., 2024), we estimated whether the copy number variation of *GA20ox1* was single or multiple by using the ratio of the average depth in the *GA20ox1* locus to the whole genome, based on the short-read resequencing data of 220 cultivars. Despite overestimation of the copy number differences, this ratio maximizes the sensitivity of finding multiple-copy-type natural variants. Homozygous multiple-copy-type cultivars such as Koshihikari (Wang *et al*., 2023; Park *et al*., 2025) and AD had more than twofold the average depth ratio of ID, TH, and TM, which have a homozygous single copy (Fig. 3a); this was reasonable considering our long-read sequencing results (Fig. 2a). These tendencies indicated that our analysis captured the differences in the ratios of average depth in the *GA20ox1* locus to the whole genome due to the copy number variation in the JAM2 founders. With a threshold of 2, the proportion of putative multiple-copy-type cultivars inside the *japonica* population was 20% (5/25) in the tropical *japonica* subgroup and 32% (40/126) in the temperate *japonica* subgroups (Fig. 3b, Table S3). With this threshold, putative multiple-copy-type cultivars were not included in the *indica* subgroup (*indica* and *aus*). These results suggested that multiple copies of *GA20ox1* are present mainly in the *japonica* subgroup, and especially in temperate *japonica* cultivars. The *ga20ox2* natural variant conserved in AD contains a 383-bp deletion in the first exon (Fig. S6a). This variant originated from the Taiwanese cultivar Dee-geo-woo-gen and is conserved in IR8, a pedigree ancestor of AD (Chandler, 1968; Ferrero-Serrano *et al*., 2019). We found that this deletion-type variant was conserved in 38 out of 220 cultivars, with the highest proportion in temperate *japonica* (24 cultivars) (Fig. 3c, Table S4). For *Hd1*, functionally strong alleles without a 36-bp insertion in the first exon were included in half (63/126) of temperate *japonica*, but they were much less common in tropical *japonica, indica*, and *aus* (9/94) (Fig. 3d, Table S5).

**Fig. 3.**
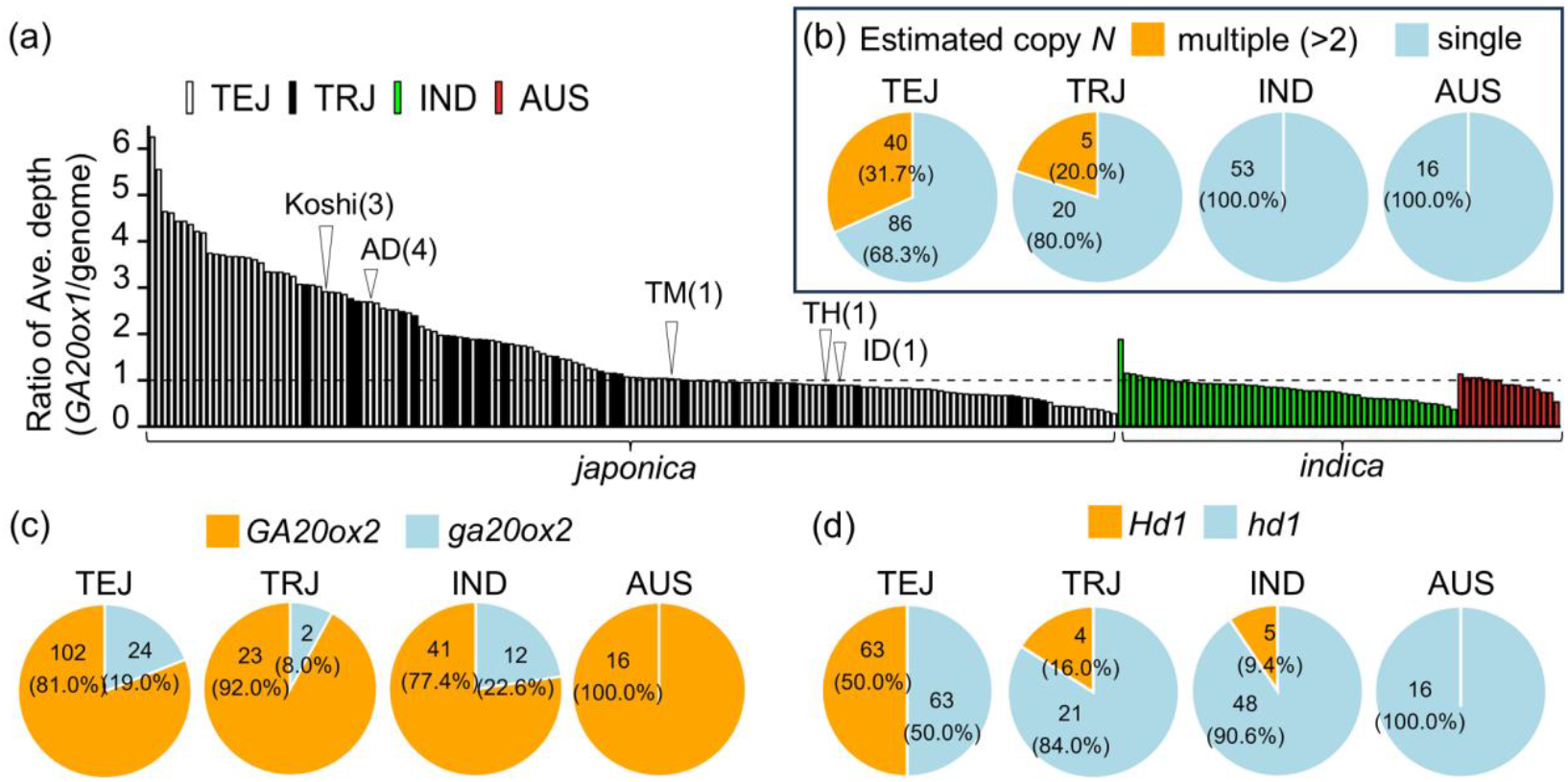
Conserved ratios of natural variations of multiple-copy *GA20ox1*, the IR8-derived deletion-type *ga20ox2* allele, and the functionally weak allele *hd1* with a 36-bp insertion in the first exon among different rice subspecies. (a) Ratio of the average depth of the *GA20ox1* locus to that of the whole genome. Numbers inside the circles represent the copy numbers identified for each JAM2 founder (in this study) and for Koshihikari (“Koshi”) in a previous study (Park *et al*., 2025). Proportions are shown of different genotypes of (b) *GA20ox1*, (c) *GA20ox2*, and (d) *Hd1*. Multiple-copy *GA20ox1* genotypes were estimated with a threshold of more than a two-fold change in the ratio of average depth (*GA20ox1* locus versus the whole genome). *ga20ox2* was marked by a >380-bp deletion in the first exon. *hd1* was marked by a 36-bp insertion in the first exon. Short-read resequencing data of 220 rice cultivars provided by the NARO (National Agriculture and Food Research Organization) Genebank (TASUKE+) were used in the analyses of the three genotypes. TEJ, temperate *japonica*; TRJ, tropical *japonica*; IND, *indica*; AUS, *aus*

These patterns indicated that there was preferential retention of non-functional *ga20ox2* and functional *Hd1* alleles in temperate *japonica* breeding compared with that in tropical regions.

### Historically selected *GA20ox1* and *GA20ox2* combinational genotypes underlying shorter CL and higher yield in elite Japanese rice cultivars

Our CH results via UAV-based time series analyses suggested that the presence of multiple copies of *GA20ox1* has the potential to alleviate the *ga20ox2*-dependent negative impacts on shoot growth at the vegetative stage (Figs. 2d, e; S6b, c). We then evaluated the combinational effects of the *GA20ox1* and *GA20ox2* genotypes on CL, shoot biomass, and yield by using the available historical field records collected between 2017 and 2021 for 52 Japanese cultivars. These cultivars’ names were registered from 1936 to 2020. Historical field record analysis showed that CL was significantly shorter in the multiple-copy *GA20ox1* and deletion-type *ga20ox2* combinational genotypes (multi/del) and the single-copy *GA20ox1* and *GA20ox2* combinational genotypes (single/non-del) than in the multiple-copy *GA20ox1* and *GA20ox2* combinational genotypes (multi/non-del) from 2017 to 2021 (Fig. 4a, upper), indicating that our dataset validated the known positive effects of *GA20ox1* and *GA20ox2* on CL (Ashikari *et al*., 2002; Sasaki *et al*., 2002; Oikawa *et al*., 2004). Shoot biomass was not changed by the variations in the combinational genotypes (Fig. 4a, middle). Notably, yield was significantly higher in the multi/del genotypes than in the multi/non-del ones in 3 of the 5 years, with a similar increasing trend observed in the remaining 2 years (Fig. 4a, lower). Such a decreasing trend in CL and an increasing trend in yield were observed from the Tohoku to the Kyushu regions of Japan (Fig. S8a,b).

**Fig 4.**
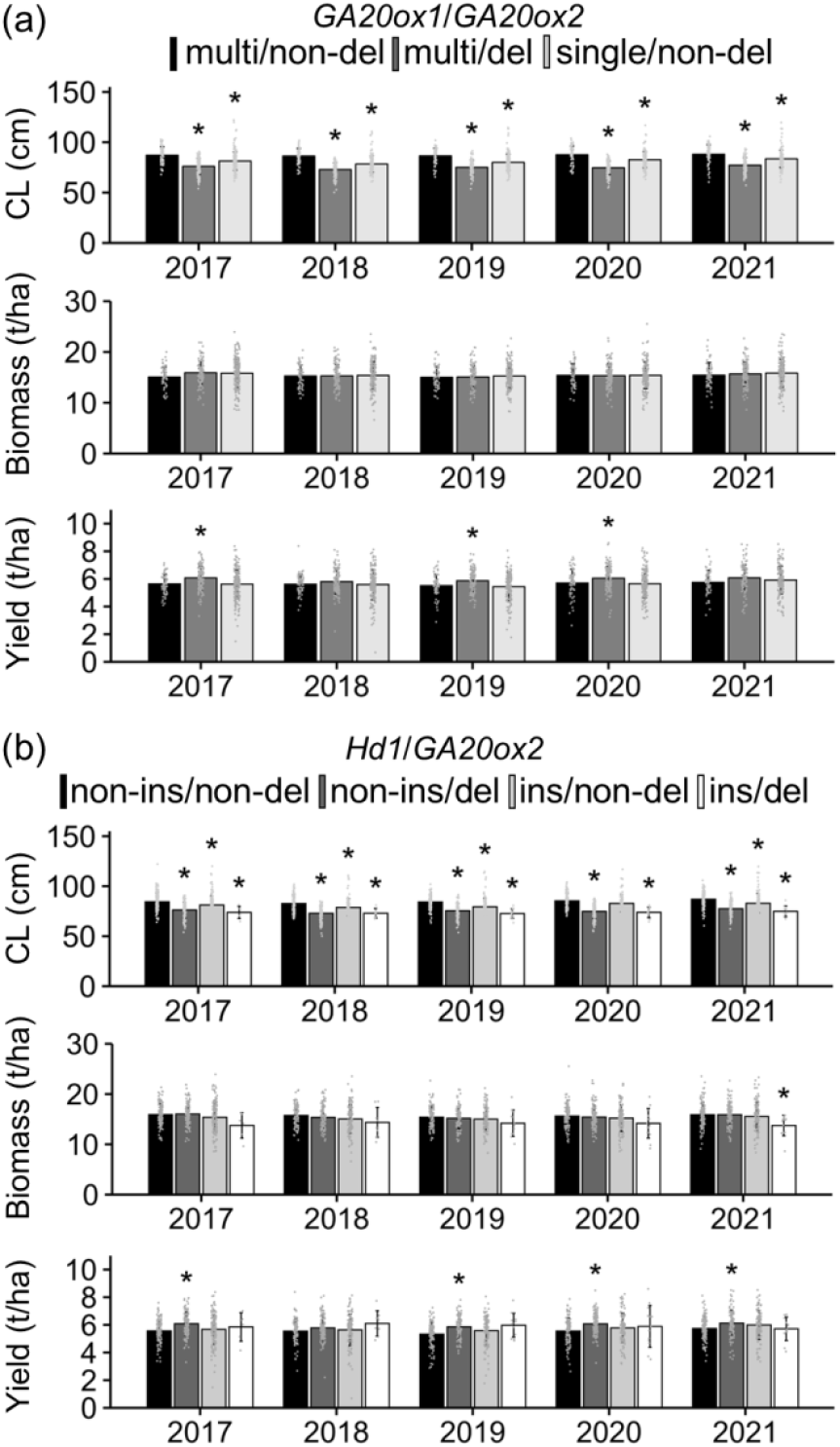
Effects of combinational genotypes of *GA20ox1, GA20ox2*, and *Hd1* on culm length, biomass, and yield in Japanese cultivars. Culm length (CL), shoot biomass, and grain yield among Japanese cultivars classified by combinational genotypes of (a) *GA20ox1* or (b) *Hd1* with *GA20ox2* under the experimental Japanese field conditions from 2017 to 2021 are shown. Each point represents the mean per cultivar × site × year for each agronomic trait, based on historical field records obtained by NARO. A total of 52 cultivars tested at up to total of 69 sites are plotted. Means ± SD are shown (CL and yield, n = (a) 58 to 206, (b) n = 6 to 147; biomass, (a) 56 to 198, (b) n = 6 to 144). Asterisks indicate significant differences from the control combinational genotypes: multiple-copy *GA20ox1* or *Hd1* allele without a 36-bp in the first exon combined with the *GA20ox2* allele without a >380-bp deletion in the first exon (i.e., combinations of each of the functionally strong genotypes) (*P* < 0.05, Dunnett’s test). multi, multiple-copy; non-del, non-deletion; del, deletion-type mutation; non-ins, non-insertion; ins, insertion-type mutation

### Historically selected *Hd1* and *GA20ox2* combinational genotypes contributing to shorter CL and higher yield

As in the multiple-copy *GA20ox1* natural variant, *Hd1* alleles without an insertion-type mutation in the first exon enhanced shoot growth (Fig. S7b,e). Thus, these functionally strong alleles may ameliorate the negative impact of *ga20ox2* on shoot biomass. Our analysis of historical field records revealed that, in at least 4 of the 5 years, when each of the genotypes was the insertion-type mutation of *hd1* (ins) or the deletion-type mutation of *ga20ox2* (del), CL was significantly shorter than that of the functionally strong combinational genotypes of *Hd1* and *GA20ox2* (non-ins/non-del) (Fig. 4b, upper). Biomass had a decreasing tendency in the insertion-type *hd1* and deletion-type *ga20ox2* combinational genotypes (ins/del) compared with the non-ins/non-del ones across the 5 years (Fig. 4b, middle). Yield was significantly greater in the non-ins/del genotypes than in the non-ins/non-del ones in 4 of the 5 years (Fig. 4b, lower). The same trends of decreased CL and increased yield were observed from the Tohoku to the Kyushu regions of Japan, as well as in the multi/del genotypes of *GA20ox1* and *GA20ox2* (Fig. S8a,b).

### Historical and geographical distribution of combinational genotypes of *GA20ox1, GA20ox2*, and *Hd1* for high-yield and shorter CL across Japanese rice-growing regions in past breeding programs

Our analysis of Japanese historical field records of elite temperate *japonica* cultivars revealed that multiple-copy *GA20ox1* and functional *Hd1* both tended to promote higher yield with *ga20ox2* among modern Japanese cultivars. This raises the question of when and how favorable combinational genotypes (multi/del/non-ins) of *GA20ox1, GA20ox2*, and *Hd1* were preserved and incorporated into Japanese breeding. To answer this, we examined the historical and geographical emergence of temperate *japonica* cultivars carrying the favorable combinational genotypes from early to modern Japanese breeding programs. Among 65 Japanese cultivars for which the breeding region and the year of name release were known, 20 showed the consistency with the presence of both the multiple-copy *GA20ox1* variant and a functionally strong *Hd1* allele, combined with the deletion-type *ga20ox2* allele (Fig. S9a). The origin of the multi/del/non-ins genotypes was traced back to the late 1980s (Fig. S9b,c). Following the release of Kinuhikari (Tabuchi *et al*., 2000), which harbors the favorable combinational genotypes of *GA20ox1, GA20ox2*, and *Hd1* (Figs. S10, S11, S12), in the Chubu region in the 1980s (Fig. S13), cultivars of the multi/del/non-ins genotypes were bred widely across rice-growing regions in Japan, particularly from the east-central to southwestern regions from the 2000s onward (Figs. 5, S14; Tables. S6, S7). Notably, among the above 65 cultivars, none of the cultivars bred in Tohoku or Hokkaido carried the deletion-type *ga20ox2* allele (Fig. S13, Table S6).

**Fig. 5.**
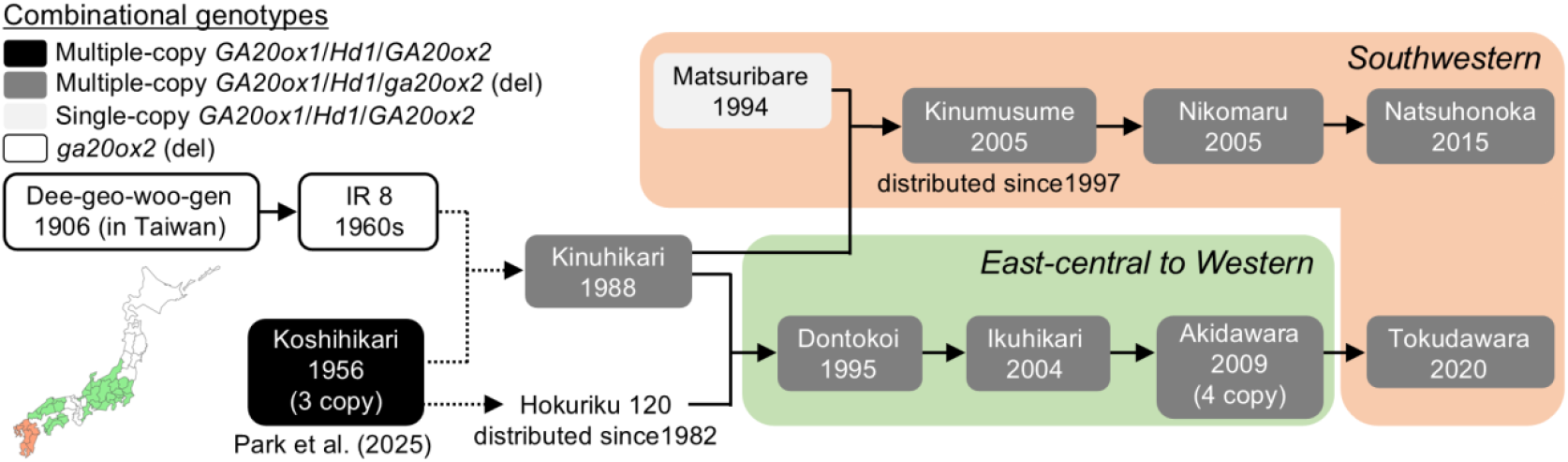
Historical emergence of combinational genotypes based on multiple-copy *GA20ox1* and functionally strong *Hd1* genotypes with the IR8-derived deletion-type *ga20ox2* allele in Japanese breeding programs. The pedigrees of Japanese cultivars harboring different combinational genotypes of *GA20ox1, Hd1*, and *GA20ox2/Sd1* are shown. These cultivars were bred across east-central to western regions (light green) or southwestern regions (light orange) of Japan from 1956 to 2020. The years below the cultivar names represent the years of name release. In this diagram, solid lines denote direct lineage connections between progeny and recipients, whereas dashed lines indicate indirect associations within the same pedigree structure. del, mutation allele with a >380-bp deletion in the first exon of *GA20ox2*

## Discussion

### Possible roles of favorable combinational genotypes of *GA20ox1, GA20ox1*, and *Hd1* in balancing shoot growth, shaping semi-dwarf architecture, and achieving high yields in modern Japanese rice-breeding programs

Dwarfing has been central to modern rice breeding for improved lodging resistance and yield, but excessive dwarfism can reduce vegetative biomass and limit yield (reviewed by Nagai *et al*., 2018; Weng *et al*., 2025). Through the characterization of identified QTLs, we found that multiple-copy *GA20ox1* genotypes positively influenced canopy height during the vegetative stage and functionally strong *Hd1* genotypes positively influenced shoot biomass (Figs. 2, S7). These functionally strong genotypes are well conserved in temperate *japonica* cultivars, in contrast to tropical *japonica* and *indica* subspecies (Fig. 3), suggesting that they play important roles in the breeding of elite temperate *japonica* cultivars. As expected, cultivars harboring both functionally strong *GA20ox1* and *Hd1* genotypes, along with the non-functional (deletion-type) *ga20ox2* allele, showed no biomass penalty under Japanese field conditions compared with those with functional *GA20ox2*, and they maintained CLs within the 60-to 80-cm range typical of modern elite Japanese cultivars (Fig. 4, Shimono *et al*., 2002). In contrast, combinations of functionally weak *hd1* and *ga20ox2* alleles resulted in reduced shoot biomass, and this may explain their limited co-conservation among elite Japanese cultivars (Fig. S13). Notably, combinational genotypes involving single-copy *GA20ox1* and *ga20ox2* were absent from our genomic dataset of 65 Japanese cultivars. These findings strongly suggest that multiple-copy *GA20ox1* and functionally strong *Hd1* genotypes are key partners with dwarfism-related variants, enabling the development of semi-dwarf cultivars without compromising shoot growth.

We next discuss how yield is influenced by the combinational genotypes of *GA20ox1, GA20ox2*, and *Hd1*. Our analysis revealed that cultivars carrying multiple-copy *GA20ox1* and a functional *Hd1* allele in combination with *ga20ox2* produced higher yields than those with *GA20ox2* (Figs. 4, S8). As reviewed by Weng *et al*. (2025), maintaining CH in short-culm ideotypes is one of the promising strategies for improving rice yield potential. This is because dwarfing raises the harvest index by allocating carbon away from elongating stems toward reproductive sinks; and traits that sustain biomass (e.g., an adequately tall canopy) preserve the spikelet sink strength (Fu *et al*., 2011; Weng *et al*., 2025). To increase canopy/plant height, multiple-copy *GA20ox1* and a functionally strong *Hd1* allele should be helpful (Fig. 2d, Fig. S7d), thereby enhancing source strength. This may be one of the reasons why these functionally strong genotypes with *ga20ox2* increase yield in a wide range of Japanese regions.

### Geographical distribution of favorable combinational genotypes for breeding semi-dwarf and high-yielding cultivars across Japanese rice-growing regions

In our 1936–2020 dataset of cultivars bred in the Hokkaido and Tohoku regions (i.e., northern Japan), no entries harbored the deletion-type *ga20ox2*/*sd1* allele that confers IR8-type semi-dwarfism and is present in AD (Figs. S6; S9a,b). This finding indicates that the *sd1*/*ga20ox2* variant derived from IR8 may have been used rarely in these regions. In our dataset, cultivars carrying the multiple-copy *GA20ox1* and deletion-type *ga20ox2* combination tended to show higher yields and shorter CLs, and this pattern held regardless of latitude (Fig. S8). These observations suggest that neither yield potential nor CL is the main constraint on adopting this favorable genotypic combination in northern breeding programs. One possible explanation is that non-functional *ga20ox2* natural variants may delay leaf development and reduce biomass accumulation during the vegetative stage under cooler, shorter growing seasons (Shimono et al., 2002). Under such conditions, breeders may have been reluctant to use dwarfism-inducing variants unless they were combined with strongly favorable partner genotypes. This context likely contributed to the low frequency of the deletion-type *ga20ox2* allele in cultivars bred in Hokkaido and Tohoku (Fig. S9a,b). Akihikari, which traces its pedigree back to Reimei (Futsuhara *et al*., 1967; Ashikari *et al*., 2002; Tomita & Ishii, 2018), provides a concrete example: despite this relationship, Akihikari does not carry the non-functional *ga20ox2* allele (Table S6). This pattern indicates that, even when the Reimei-derived *ga20ox2*/*sd1* allele was available in northern breeding populations, lines retaining it were not favored, probably because it impaired vegetative growth under the prevailing cool conditions.

Under these constraints on *ga20ox2*/*sd1* use, alleles that enhance yield without strongly penalizing growth in cool temperate climates become particularly important. Multiple-copy *GA20ox1* has also been shown to increase yield in many modern Chinese *japonica*/*geng* cultivars, particularly in Liaoning Province, which lies at a latitude similar to Tohoku and Hokkaido (Wang et al., 2023). This independent evidence supports the idea that the *GA20ox1* tandem repeat can enhance yield potential even in cooler temperate rice-growing regions. Overall, our results suggest that the tandem repeat natural variant of *GA20ox1* is a useful molecular target for marker-assisted selection (reviewed by Jena & Mackill, 2008). With the markers developed in this study (Fig. S10; Table S2), breeders can combine this variant with *ga20ox2* to develop semi-dwarf, high-yielding cultivars that perform well even under cool conditions in temperate rice-growing regions of Asia.

## Supporting information

Supplementary_Documentation

## Acknowledgements

We thank Makiko Suzuki and Yuko Aono for their technical and field support. We appreciate the technical staff of the Institute of Crop Science, NARO, for their help in managing the rice fields. We would like to thank Dr. Itoh for providing the NIL harboring *hd1* allele. We are grateful to Dr. Goto and Dr. Matsushita for sharing historical field records and seeds of Japanese cultivars. We thank the researchers of NARO and the Japanese Public Research Institute in every prefecture involved in the collection of historical field records. We thank Haruki Nakamura for his guidance on how to analyze Nanopore sequencing data. This research was supported by the research program from the Ministry of Agriculture, Forestry and Fisheries of Japan [grant number: DIT2001 and J012037] and by JSPS Grant-in-Aid for Transformative Research Areas (A) (23H04756 and 23H04968). This research was partially supported by the Research Supporting Program of the Research Center for Advanced Analysis, NARO. We are thankful to the editors at ELSS, Inc. (https://elss.co.jp/en/) for their professional editing services before submission.

## Author contributions

**HF:** Writing – original draft and review & editing, Data acquisition, Data curation, Data analysis, Visualization, Conceptualization. **AF:** Data acquisition, Data curation, Data analysis. **TS:** Data acquisition, Data curation. **YK:** Data acquisition, Data curation, Data analysis. **KN:** Data acquisition, Data curation. **TK:** Data acquisition. **JY:** Writing – review & editing, Data acquisition, Data curation, Data analysis, Funding acquisition, Conceptualization, Supervision. **DO:** Writing – review & editing, Data acquisition, Data curation, Data analysis, Visualization, Conceptualization, Supervision.

## Data availability

The datasets used and/or analyzed in the current study are available from the corresponding author upon reasonable request.

