## Supplementary_Figs for "Dissecting Agronomically Favorable Genotypes in Temperate Japonica Rice via Haplotype Analysis of a Japan-MAGIC Population"

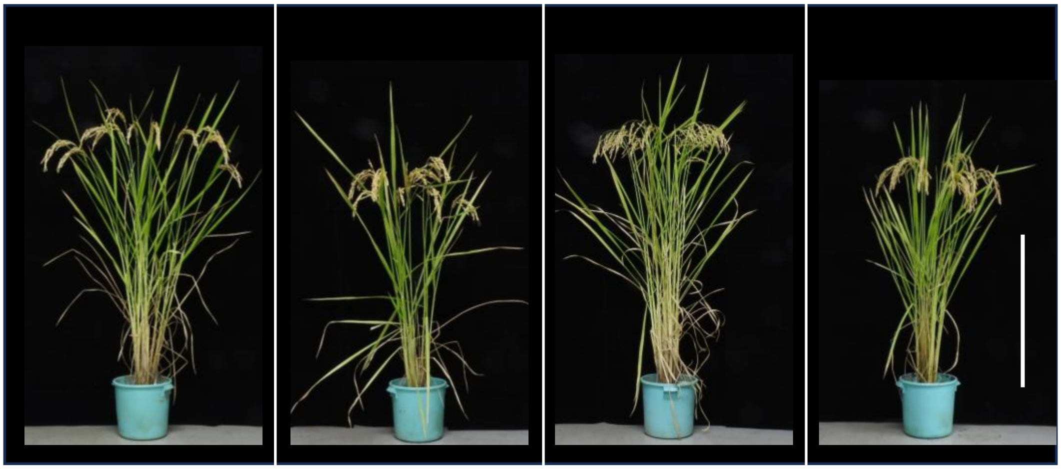

Akidawara (AD) | Iwaidawara (ID) | Tachiharuka (TH) | Toyomeki (TM)

**Fig. S1. JAPAN-MAGIC2 founders.** Founders of JAPAN-MAGIC2 (JAM2). A rice plant of each cultivar was brought in from the paddy field at the ripening stage and photographed. White bar, 50 cm.

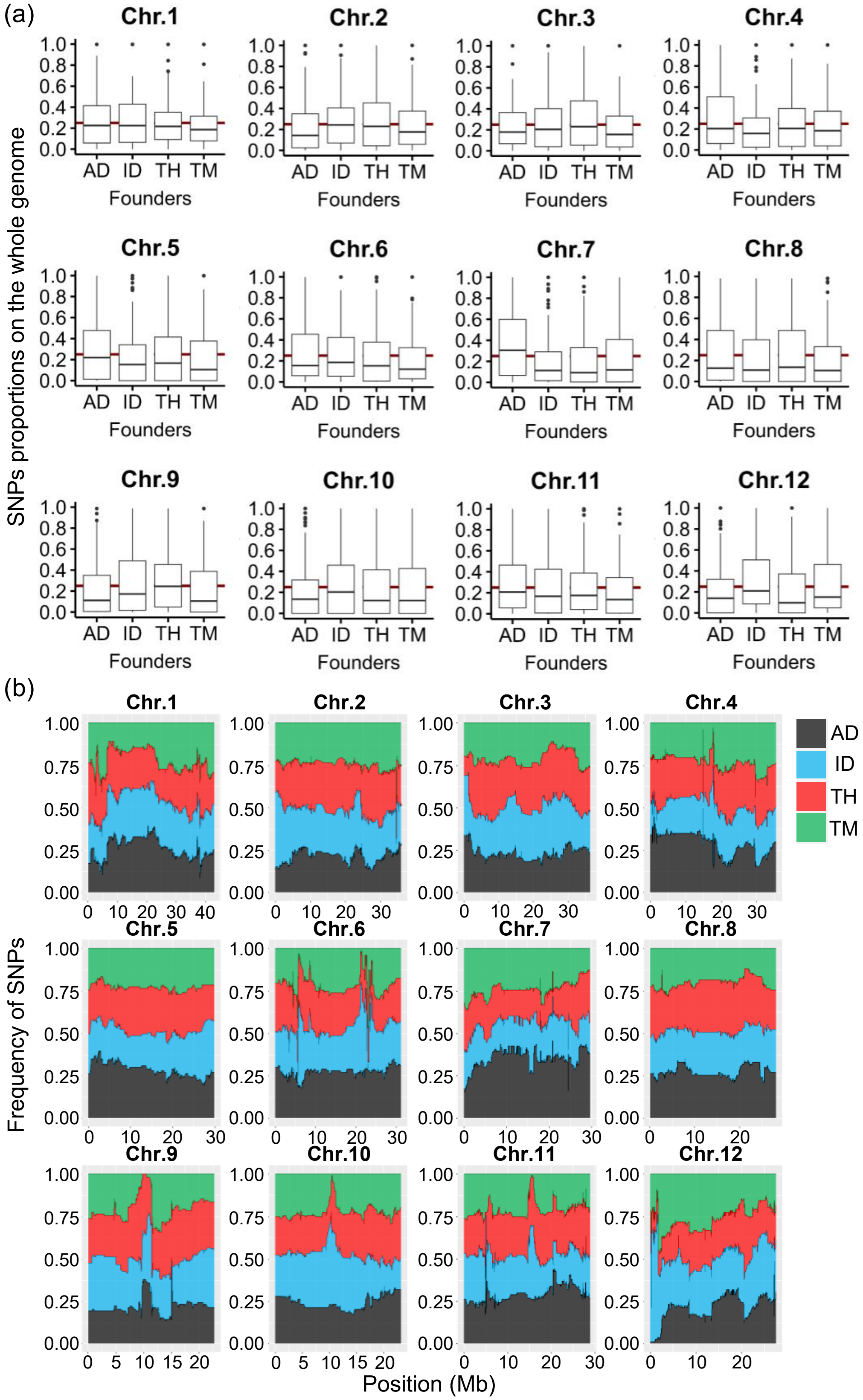


**Fig. S2. Distribution of single nucleotide polymorphism in each chromosome of JAM2 lines.** (a) Proportions of founder-derived single nucleotide polymorphisms (SNPs) across chromosomes 1 to 12 in JAM2 lines. Founders are categorized as Akidawara (AD), Iwaidawara (ID), Tachiharuka (TH), and Toyomeki (TM). (b) Frequency of SNPs along chromosomes 1 to 12. Frequencies are color coded: AD (black), ID (blue), TH (red), and TM (green).

(a)

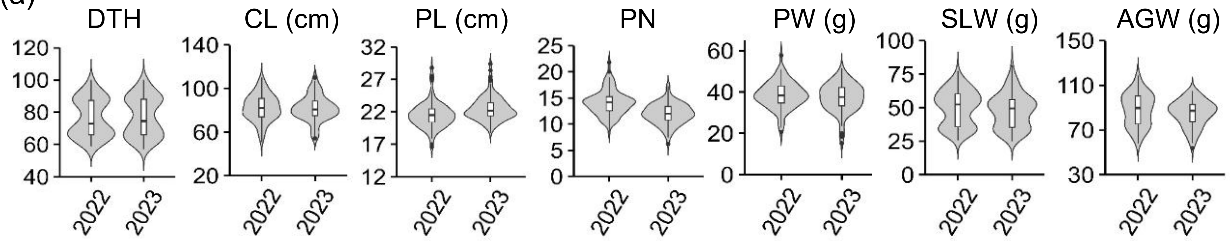


(b)

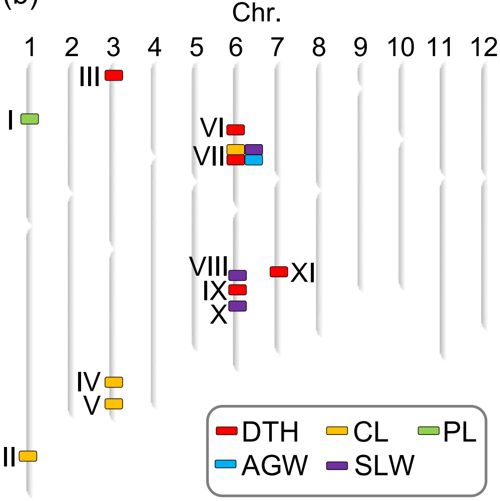


(c)

| (QTL) No | Chr | Trait | Year 2022 AD | Year 2022 ID | Year 2022 TH | Year 2022 TM | Year 2023 AD | Year 2023 ID | Year 2023 TH | Year 2023 TM |
|---|---|---|---|---|---|---|---|---|---|---|
| I | 1 | PL | 21 | 23.1 | 20.8 | 21 | 21.9 | 24 | 21.6 | 22.3 |
| II | 1 | CL | 67.8 | 85.6 | 85 | 84.3 | 65 | 84.9 | 84.2 | 85 |
| III | 3 | DTH | 83.2 | 77.5 | 75.7 | 67.6 | 84.3 | 78.9 | 75.8 | 68.3 |
| IV | 3 | CL | 91.1 | 81.3 | 80.1 | 77.8 | 91 | 78.1 | 79.2 | 79 |
| V | 3 | CL | 91.1 | 77.2 | 79.7 | 80.4 | 91.8 | 76.3 | 78.6 | 78.9 |
| VI | 6 | DTH | 84.7 | 68.2 | 83 | 69.6 | 86.3 | 68.5 | 84 | 69.7 |
| VII | 6 | CL | 88.6 | 74.3 | 85.6 | 79.8 | 87.5 | 75.2 | 85.8 | 76.5 |
| VII | 6 | SLW | 58.1 | 37.7 | 59.4 | 40.4 | 54.7 | 37.3 | 57.7 | 37.8 |
| VII | 6 | DTH | 85.5 | 65.8 | 83.9 | 66.9 | 87.1 | 65.9 | 84.9 | 66.9 |
| VII | 6 | AGW | 96.2 | 72.8 | 95.7 | 77.9 | 91.3 | 76.5 | 91.9 | 77.2 |
| VIII | 6 | SLW | 55.8 | 43.6 | 57 | 54.3 | 52.2 | 42 | 55.9 | 52.7 |
| IX | 6 | DTH | 80.8 | 84.6 | 82.9 | 70.4 | 81.8 | 83.8 | 84.7 | 70.8 |
| X | 6 | SLW | 91 | 80.6 | 92.5 | 91.4 | 88.8 | 80.1 | 90 | 87.4 |
| XI | 7 | DTH | 72.4 | 80.4 | 72 | 83.8 | 72.5 | 82.2 | 73 | 84.1 |

**Fig. S3. Putative QTLs associated with agronomic traits conserved in JAM2 founders.** (a) Distributed values of observed agronomic traits across JAM2 lines in 2022 and 2023 (n = 100). (b) Map of putative QTLs determined in this study. Each roman numeral represents a locus in Table 1. (c) Mean values of each trait calculated after classification by using the genomic regions of putative QTLs. These regions are shown in Table 1. PL, panicle length; CL, culm length; DTH, days to heading; SLW, stem and leaf dry weight; AGW, aboveground dry weight

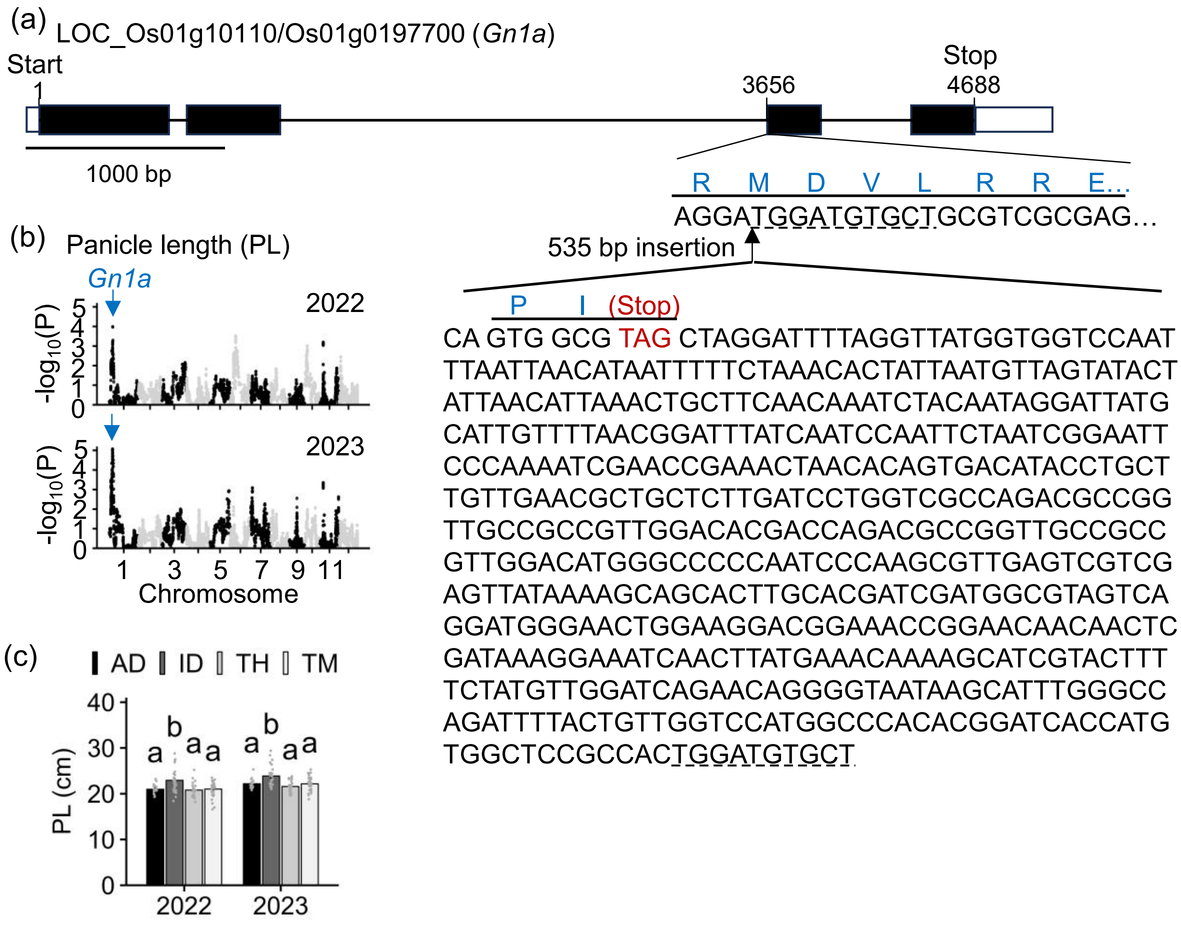


**Fig. S4. Gene structure of an insertion-type *gn1a* variant conserved in Iwaidawara.** (a) Scheme of novel natural variant of *Gn1a*. Black boxes indicate the exons, white boxes indicate untranslated regions, and black lines connecting black boxes indicate the introns. The inserted sequence of 535 bp and predicted amino-acid sequence (blue letters) in the third exon of the Iwaidawara allele are shown. (b) Genome-wide association study detecting the peak including the *Gn1a* locus in the JAM2 population in 2022 and 2023. Each plot represents a single nucleotide polymorphism. (c) Panicle length (PL) among JAM2 lines classified by the haplotype including the *Gn1a* locus. Means ± SD are shown (n = 16 to 33). Different letters indicate significant differences among JAM2 lines ($P < 0.05$, Tukey–Kramer test). AD, Akidawara; ID, Iwaidawara; TH, Tachiharuka; TM, Toyomeki.

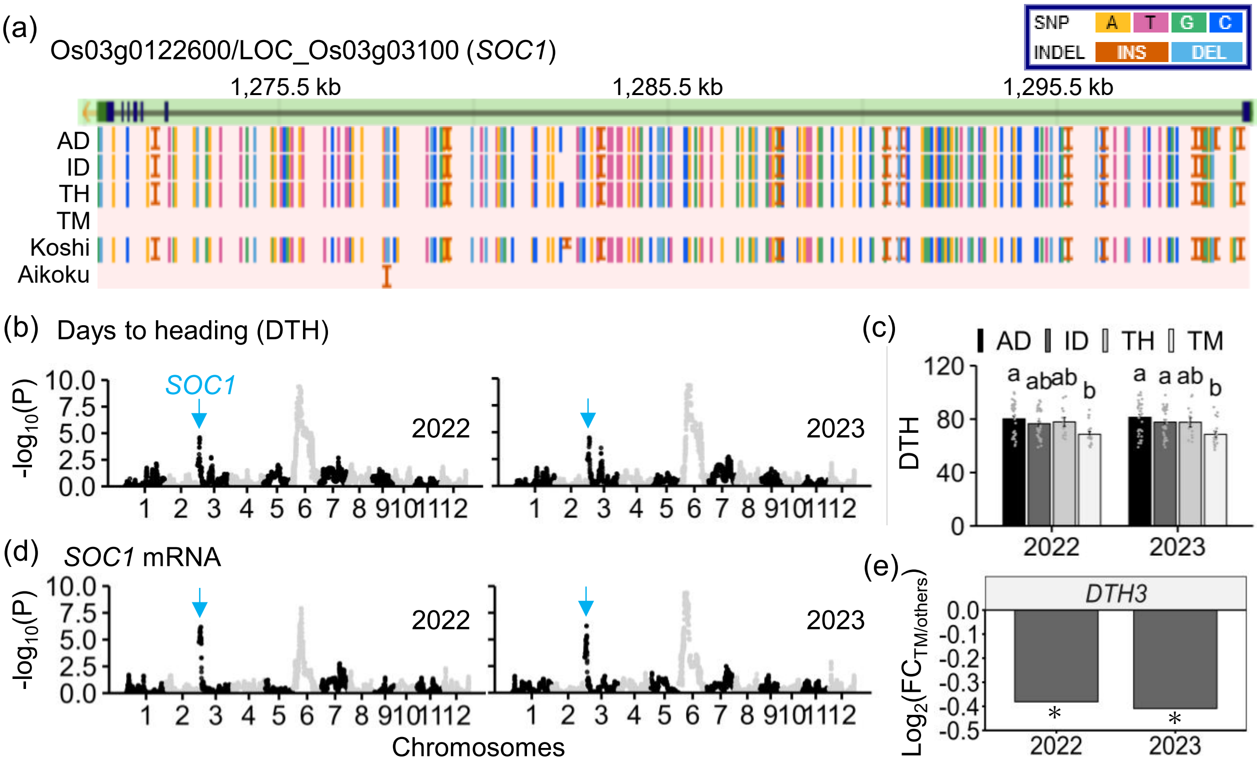


**Fig. S5. Effect of the putative QTL corresponding to *SOC1* on days to heading.** (a) Visualization of *SOC1* variants among the four founders, Koshihikari ('Koshi'), and Aikoku. Polymorphisms and indels relative to Nipponbare are highlighted using different colors and symbols. (b) Genome-wide association study (GWAS) for days to heading (DTH) detecting the peak including the *SOC1* locus among JAM lines. (c) DTH among JAM2 lines classified by haplotypes including the *SOC1* locus. Means ± SD are shown (n = 12 to 36). Different letters indicate significant differences among genotypes ($P < 0.05$, Tukey–Kramer test). (d) Expression GWAS of *SOC1* among JAM2 lines. (e) Differences in the mRNA levels of *SOC1* between lines harboring the Toyomeki-derived *soc1* allele and those harboring the other founder-derived *SOC1* alleles. Asterisks indicate significant differences between the haplotypes of Toyomeki and the other founders among the JAM2 lines (padj < 0.05, n = 19 to 81). AD, Akidawara; ID, Iwaidawara; TH, Tachiharuka; TM, Toyomeki, FC, fold change.

(a)

LOC_Os01g66100/Os01g0883800 (*GA20ox2*)

| Variety | Genotype | Deletion position<br>From 1st to 2nd exon | Origin | Reference |
|---|---|---|---|---|
| ID / TH / TM | *Sd1* | - | - | - |
| AD | *sd1* | -383 bp | Dee-geo-woo-gen | Murai et al., 2004 |

(b) (c)

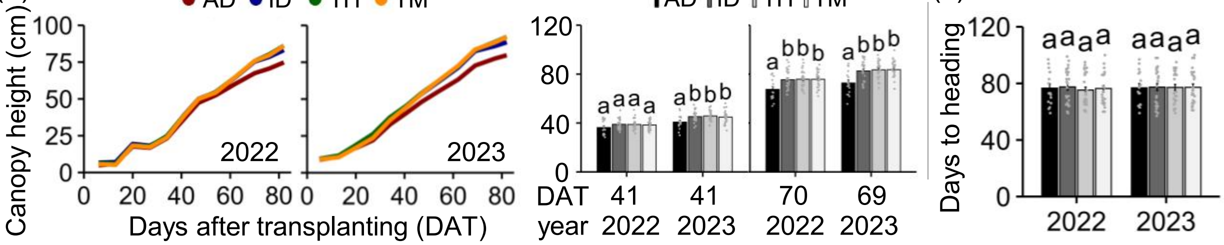


**Fig. S6. Time series of canopy height at the vegetative stage in JAM2 lines classified by the haplotype including the *GA20ox2* locus.** (a) Variation of *GA20ox2* genotypes derived from Akidawara (AD), Iwaidawara (ID), Tachiharuka (TH), and Toyomeki (TM). Time series from the middle of May to late August of (b) canopy height (CH, cm) and (c) days to heading (DTH) in JAM2 lines classified by the haplotype including the *GA20ox2* locus. Plants were transplanted to experimental fields on 18 May 2022 (0 DAT) and 17 May 2023 (0 DAT). Means ± SD for CH at two time points, and DTH (n = 23 to 26), are shown. Time series of CH are shown with means of each group classified by *GA20ox2* genotypes derived from each JAM2 founder.

(a) LOC_Os06g16370/Os06g0275000

| Variety | Genotype | Indels position at 1st exon | | | Alleles in reference | Reference |
|---|---|---|---|---|---|---|
| | | 327 | 468–500 | 702–744 | | |
| AD/TH/Koshi | *Hd1* | - | | | Nipponbare | |
| TM | *hd1/se1* | +36 bp | | | Ginbouszu | Yano et al., 2000 |
| ID | *hd1/se1* | +36 bp | | -43 bp | HS66 | Yano et al., 2000 |
| Kasa | *hd1* | +36 bp | -33 bp | | Kasalath | Yano et al., 2000 |

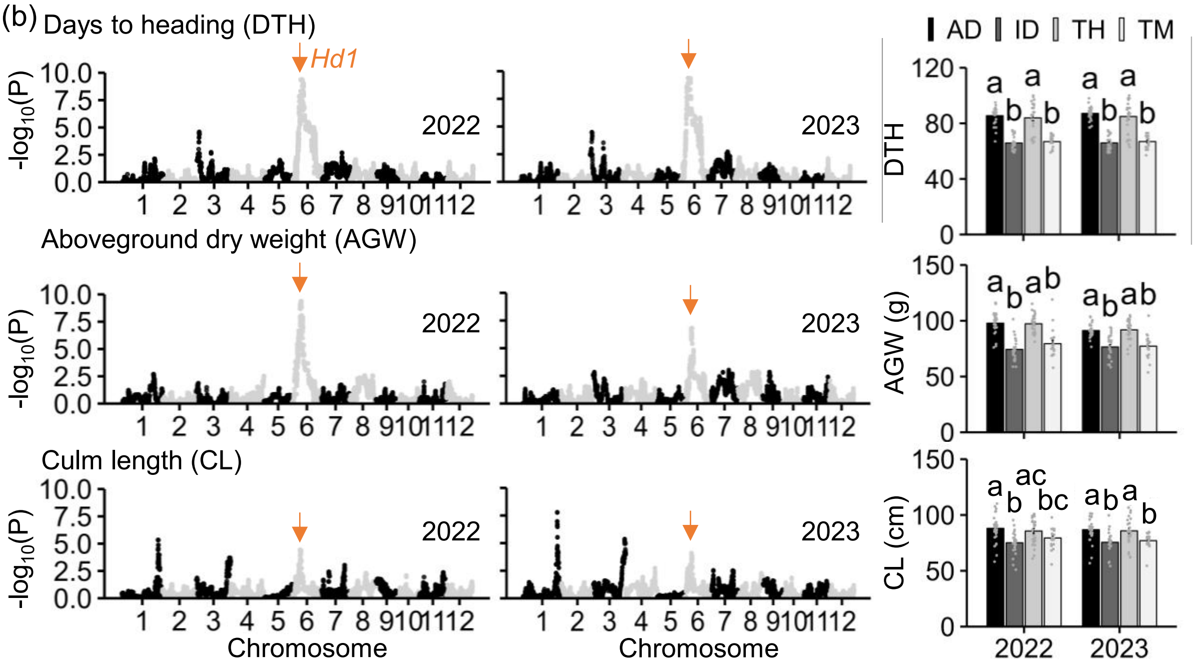


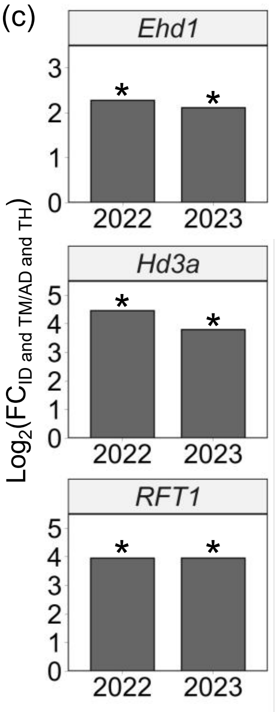


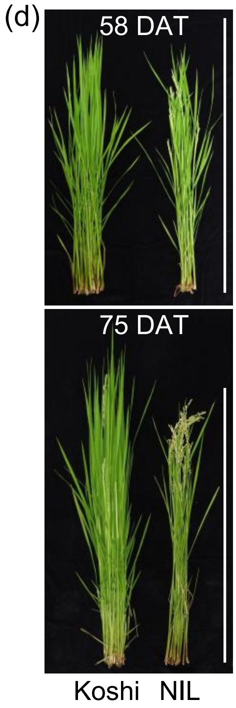


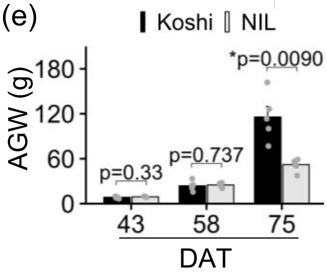


**Fig. S7. Effects of the putative QTL containing *Hd1* on days to heading, culm length, canopy height at the vegetative stage, and shoot biomass.** (a) Variation of *Hd1* genotypes derived from Akidawara (AD), Iwaidawara (ID), Tachiharuka (TH), Toyomeki (TM), Koshihikari (Koshi), and Kasalath (Kasa). (b) Days to heading (DTH), aboveground weight (AGW), and culm length (CL) among JAM2 lines classified by haplotypes including the *Hd1* locus. Means ± SD are shown (n = 20 to 28). Different letters indicate significant differences among cultivars ($P < 0.05$, Tukey–Kramer test). (c) mRNA levels of positive flowering regulators in JAM2 lines: *Early heading date 1* (*Ehd1*), *heading date 3a* (*Hd3a*), and *RICE FLOWERING LOCUS T 1* (*RFT1*). Means computed depending on the haplotype groups are shown (n = 20 to 28). Asterisks indicate significant differences between the haplotypes of ID and TM vs. AD and TH among JAM2 lines (padj < 0.05, n = 45 to 55) (d) Rice plants and (e) time series of AGW in Koshi and an *Hd1* near-isogenic line (NIL) grown under experimental field conditions in 2023. The NIL harbors an insertion-type mutation of the *hd1* allele in the Koshi genetic background. NIL plants headed at 58 days after transplanting (DAT) and the Koshi plants headed at 75 DAT. Means ± SD are shown (n = 5). Asterisks and numbers above each bar indicate significant differences ($P < 0.05$, Student's *t*-test) and *P*-values, respectively. Scale bars, 100 cm.

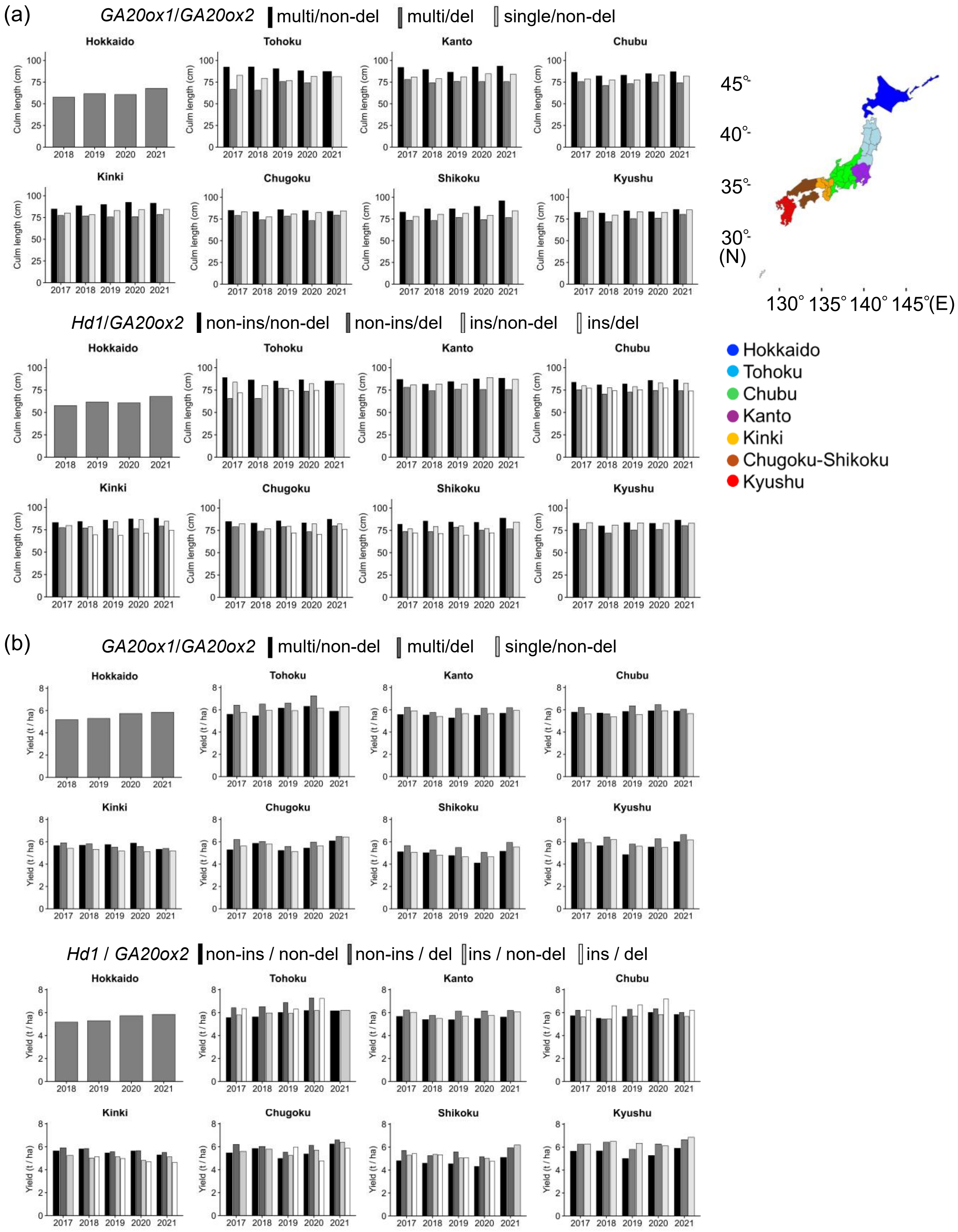


**Fig. S8. Culm length and yield affected by combinational genotypes of *GA20ox1*, *Hd1*, and *GA20ox2* in experimental fields from 2017 to 2021.** (a) Culm length and (b) yield of 52 cultivars, classified by *GA20ox1*, *Hd1*, and *GA20ox2* combinational genotypes. The results in these cultivars are consistent with the analysis shown in Fig. 4. Means in each region per year are shown in a combinational genotype–dependent manner. multi, multiple-copy; non-del, non-deletion; del, deletion-type mutation; non-ins, non-insertion; ins, insertion-type mutation

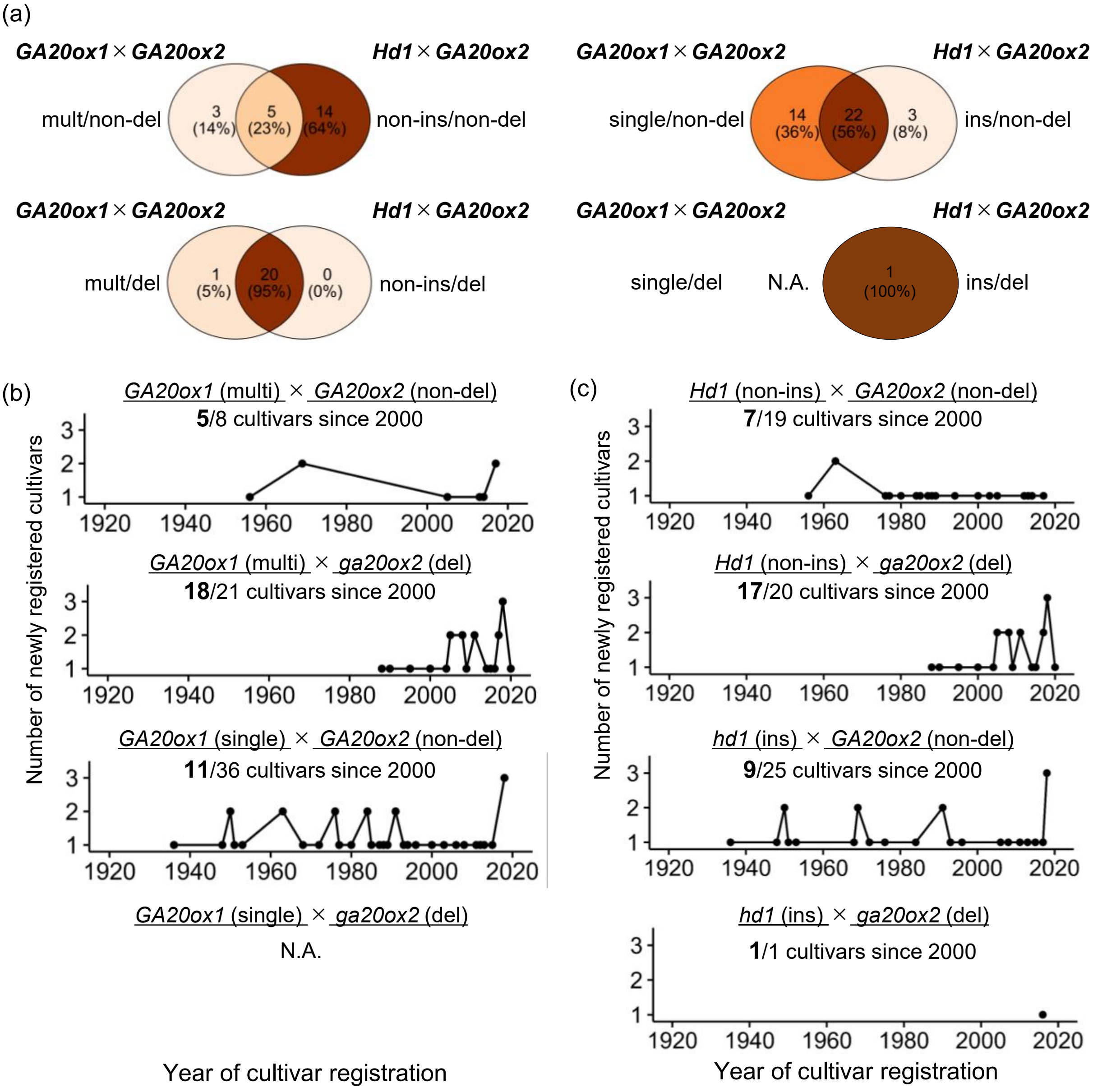


**Fig. S9. Historical emergence of combinational genotypes of *GA20ox1* and *GA20ox2*, and of *Hd1* and *GA20ox2*.** (a) Venn diagram of 65 Japanese cultivars classified according to *GA20ox1*–*GA20ox2* combinational genotypes or *Hd1*–*GA20ox2* combinations. multi, multiple-copy; non-del, non-deletion; del, deletion. Historical emergence of combinational genotypes of (b) *GA20ox1* and (c) *Hd1* with *GA20ox2* among 65 elite temperate *japonica* cultivars bred in Japan. These cultivars are the same as those used in Fig. 4c.

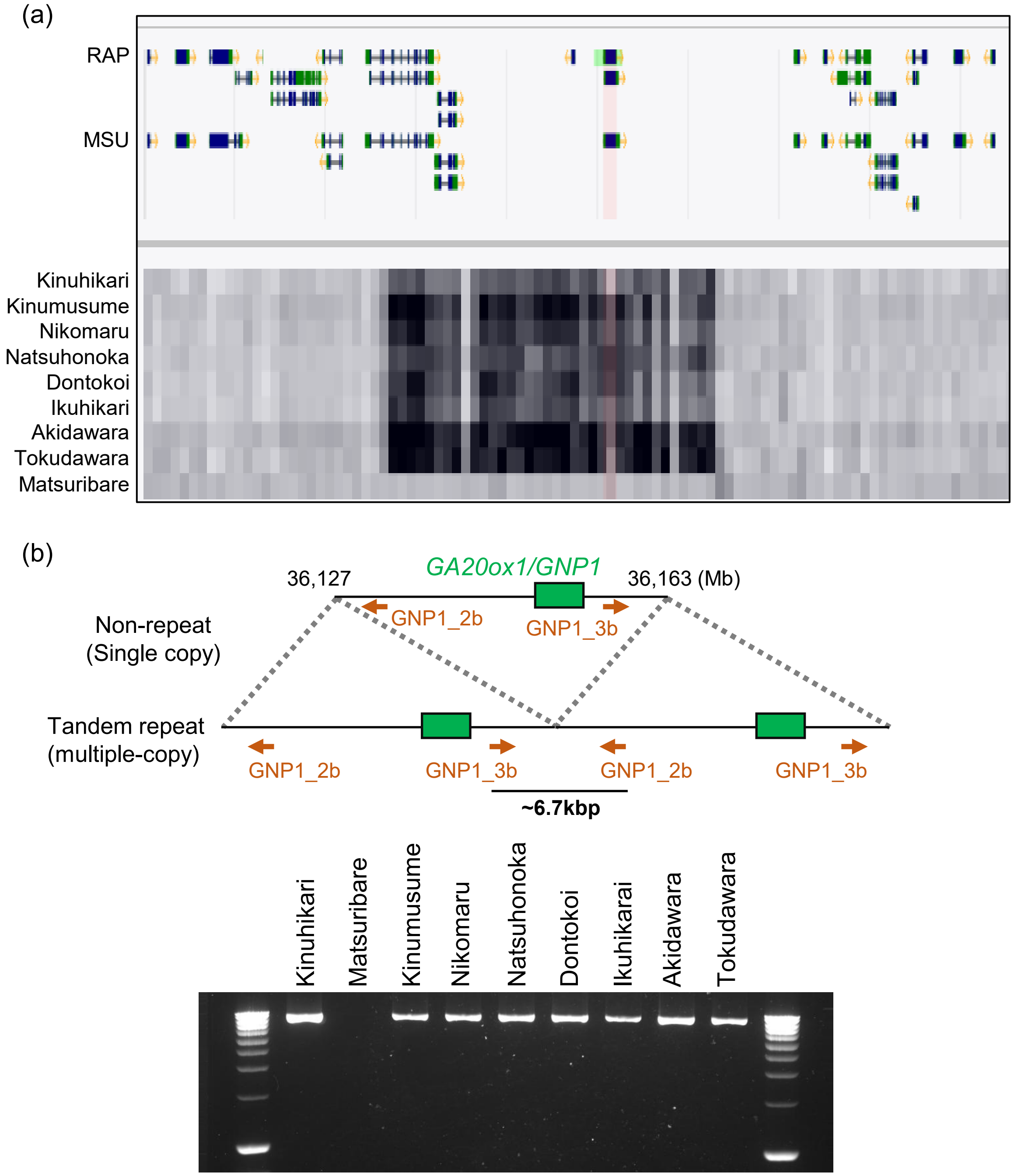


**Fig. S10. PCR-based assay for detecting duplicated region of *GA20ox1* locus.** (a) Window of the TASUKE+ browser (https://tasuke.dna.affrc.go.jp/) illustrating an estimated tandem repeat region including the genomic *GA20ox1* locus (Os03g0856700/LOC_Os03g63970) in the Japanese cultivars shown in Fig. 5. The variation in read depth for each accession is represented by the black color gradient shown in the lower panel just below the center. An approximately 36-kb genomic region from 36,127 kb to 36,163 kb shows higher read depth in cultivars harboring the estimated strong-type *GA20ox1* genotype with a threshold of 2 × the average depth ratio (locus/the whole genome) and in Kinuhikari compared with Matsuribare (at the lowest position), suggesting the presence of a tandem repeat region in Kinuhikari. Pink highlighting from the upper to the lower panel represents the coding region of the Os03t0856700-01 transcript. (b) PCR-based assay to detect non-repeat and tandem-repeat genomic regions including the *GA20ox1* locus. Primer positions for distinguishing between different alleles are shown in orange.

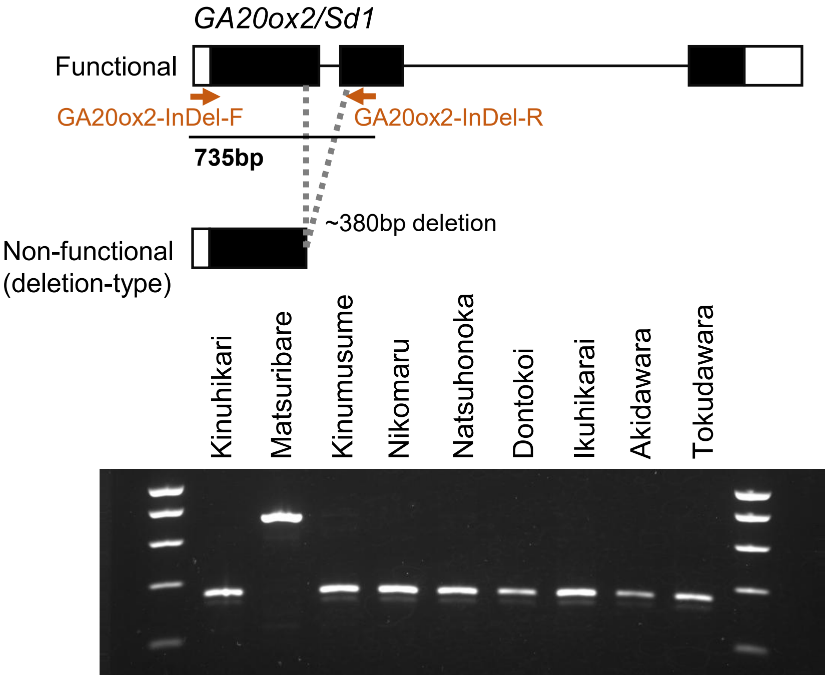


**Fig. S11. PCR-based assay to detect deletion-type mutation of the *GA20ox2/Sd1* allele.** PCR products are shown distinguishing *GA20ox2* (Os01g0883800/LOC_Os01g66100) alleles in the Japanese cultivars shown in Fig. 5. Black boxes indicate the exons, white boxes indicate untranslated regions, and black lines connecting black boxes indicate the introns. Primer positions for distinguishing between different alleles are shown in orange.

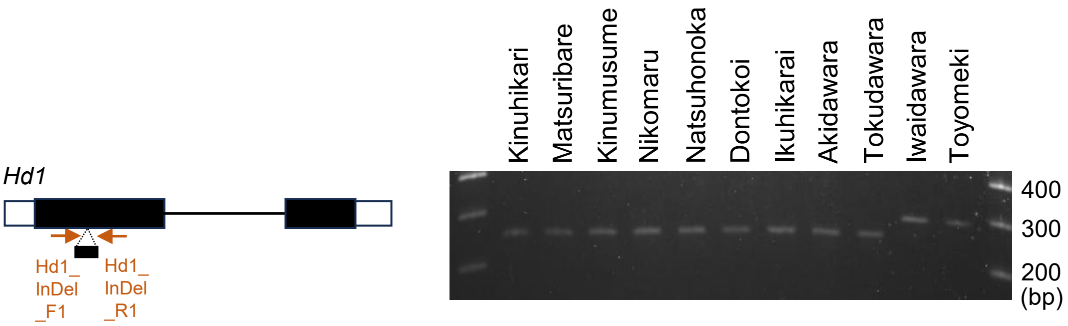


**Fig. S12. PCR-based assay to detect non-functional *hd1* natural variants.** PCR products are shown distinguishing *Hd1* (Os06g0275000/LOC_Os06g16370) alleles in the Japanese cultivars possessing the combinational genotypes of multiple-copy *GA20ox1* and non-functional *ga20ox2* (Fig. 5). Black boxes indicate the exons, white boxes indicate untranslated regions, and black lines connecting black boxes indicate the introns. Iwaidawara and Toyomeki are control cultivars harboring *hd1* variants with a 36-bp insertion in the first exon. Primer positions used to distinguish between different alleles are shown in orange.

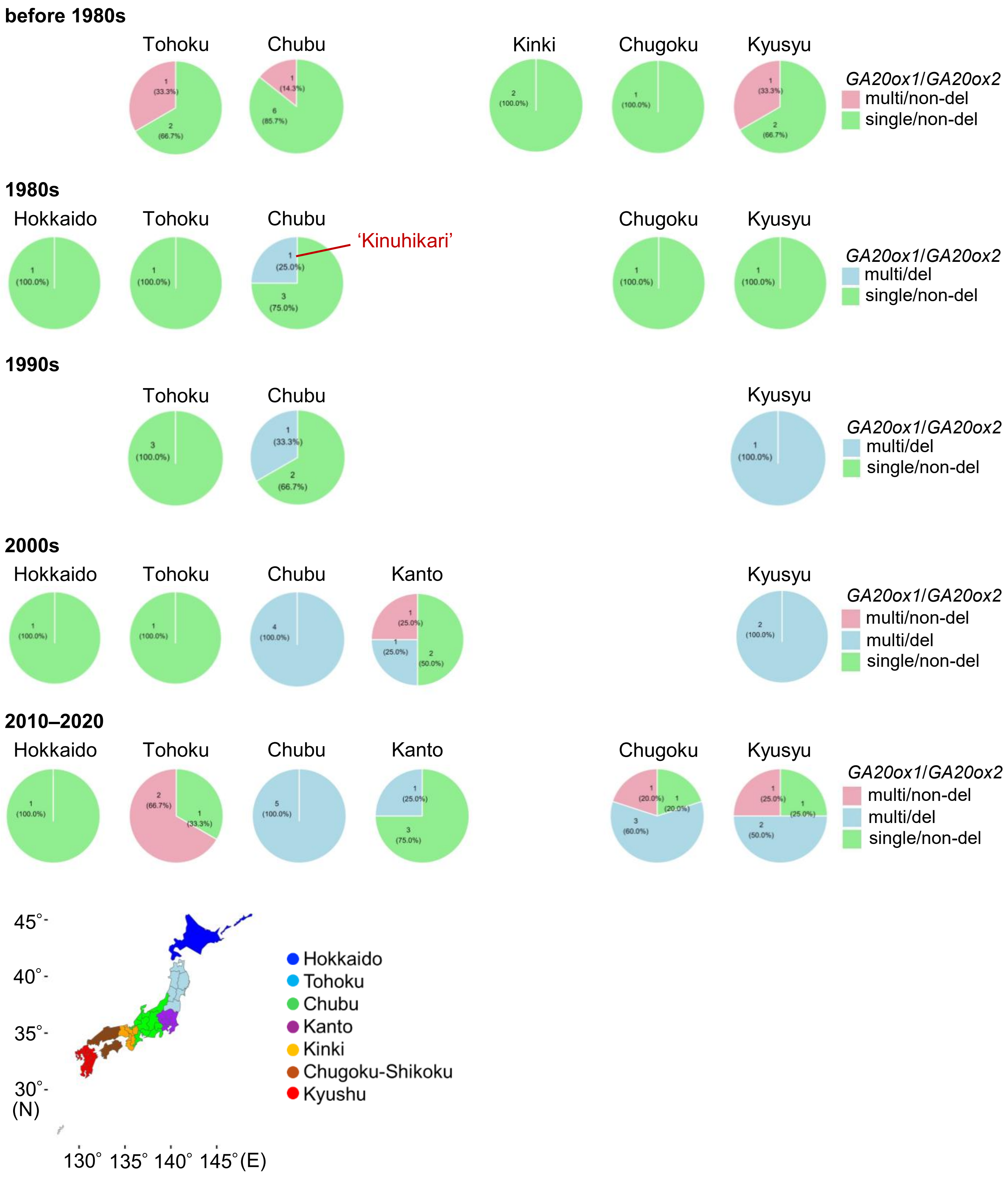


**Fig. S13. Distribution of combinational genotypes of *GA20ox1* and *GA20ox2* in different Japanese regions at different ages.** 65 Japanese cultivars, classified by *GA20ox1* and *GA20ox2* combinational genotypes, are the same as those shown in Fig. S9. multi, multiple-copy; non-del, non-deletion; del, deletion

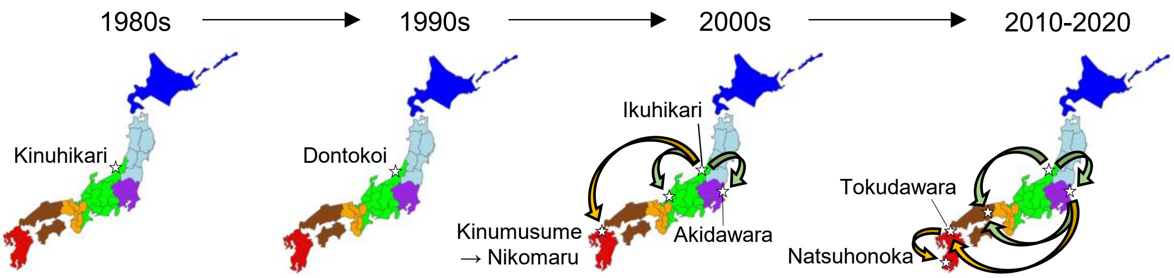


**Fig. S14. Japanese breeding history of the development of rice cultivars harboring the multiple-copy *GA20ox1* and functional *Hd1* with the IR8-derived non-functional *ga20ox2* combinational genotype.** Geographical distribution of Japanese cultivars harboring the favorable combinational genotypes for high yield and shorter culm length across different Japanese regions from the 1980s to 2020. Favorable combinational genotypes are defined as a set of multiple-copy *GA20ox1*, functionally strong *Hd1* alleles without a 36-bp insertion in the first exon, and *ga20ox2* alleles with a >380-bp deletion in the first exon. Each colored area denotes a region of Japan, ordered from north (Hokkaido) to south (Kyushu). Delineated areas within each region correspond to individual prefectures. Light green arrows indicate the distribution direction of favorable combinational genotypes from the east-central to western regions of Japan, whereas orange arrows represent their spread toward the southwestern regions.
