## Supplementary_Documentation for "Dissecting Agronomically Favorable Genotypes in Temperate Japonica Rice via Haplotype Analysis of a Japan-MAGIC Population"

**Results and Discussion: *Grain number 1a* (*Gn1a*)**

A PCR-based assay detected a band corresponding to the insertion-type *gn1a*$^{\mathrm{ID}}$ variant in Bekogonomi, a cultivar that is used for feed and has the same pedigree as Iwaidawara (ID) (Supplementary documentation Fig. 1). The PCR products from Fukuhibiki, one of the parental cultivars of Bekogonomi, indicated the presence of a non-insertion-type *Gn1a* allele, suggesting that the *gn1a* variant in ID was derived from the high-yielding line 97UK-46. The fact that this novel variant was derived from a cultivar bred for livestock use in Japan indicates that it has rarely been conserved among table Japanese rice cultivars.

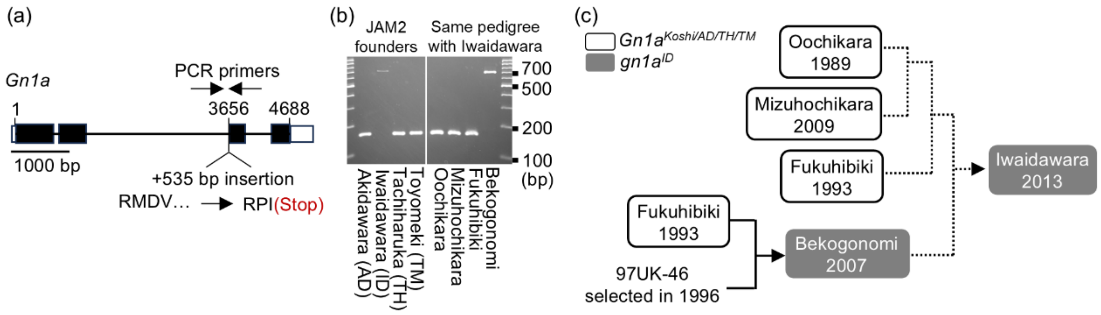


**Supplementary documentation Fig. 1. Gene structure and origin of the Iwaidawara *Gn1a* allele.** (a) Gene structure of *Gn1a* (LOC_Os01g10110/Os01g0197700) in Iwaidawara (ID). Black boxes indicate the exons, white boxes indicate untranslated regions, and black lines connecting black boxes indicate the introns. The length of the inserted sequence is 535 bp. Predicted amino-acid sequence (black letters) in the third exon of the ID allele is changed from RMDV to RPI(Stop) relative to the Nipponbare allele. (b) PCR fragments of the third exon of *Gn1a*. Primer position is described in panel a. (c) Pedigree of Japanese cultivars possessing the *gn1a*$^{\mathrm{ID}}$ variant. Solid lines denote direct lineage connections between progeny and recipients, whereas dashed lines indicate indirect associations within the same pedigree structure.

**Results and Discussion:** ***Suppressor of overexpression of constans 1*** **(*SOC1*) and *Heading date 1* (*Hd1*)**

By analyzing short-read resequencing data from 182 cultivars (6 wild, 72 landrace, and 112 cultivated rice) in TASUKE+, the multiple genome browser of the Rice Annotation Project Database for the NARO Genebank World Rice Core Collection, we found that 4.4% (8/182) of cultivars harbored the *SOC1*$^{\mathrm{Nipponbare\ (Nip)}}$ allele, which was confined to temperate *japonica* (Supplementary documentation Fig. 2a). A combinational analysis of historical data on cultivar registration (i.e., name release) and genomic information revealed that the origin of *SOC1*$^{\mathrm{Nip}}$ and *SOC1*$^{\mathrm{Koshihikari\ (Koshi)}}$ could date back to the Meiji era (Supplementary documentation Fig. 2b). Aikoku—the possible origin of *SOC1*$^{\mathrm{Nip/TM}}$—was bred in Miyagi Prefecture and grown as one of the three major cultivars from the Meiji era to the early Shōwa period. Aikoku was spread across the southern parts of the Tohoku region to Aichi Prefecture in the Chubu region, where Nipponbare was bred in 1963. *SOC1*$^{\mathrm{Nip}}$ was inherited by cultivars developed in Aichi and Fukui Prefectures and was subsequently introduced into Toyomeki, which was bred in Ibaraki Prefecture and released in 2015. As Koshihikari has been a primary cultivar in rice-growing regions of Japan since the 1970s, most Japanese cultivars including Akidawara, Iwaidawara, and Tachiharuka would harbor *SOC1*$^{\mathrm{Koshi}}$.

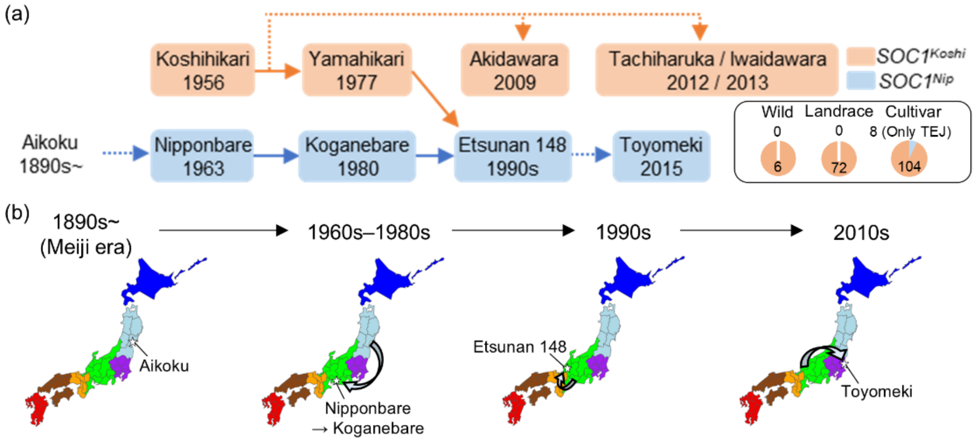


**Supplementary documentation Fig. 2. Origin of the Toyomeki *SOC1* allele.** (a) Visualization of the pedigree of Japanese cultivars harboring different *SOC1* (LOC_Os03g03100/Os03g0122600) variants derived from Koshihikari (Koshi) (orange boxes) and Nipponbare (Nip) (light blue boxes). Each year represents the year of name release for each cultivar. Short-read resequencing data of 182 cultivars were used to display the proportions of each *SOC1* variant among three categories: wild, landrace, and cultivated rice. In this diagram, direct (solid lines) and indirect (dashed lines) lineage connections between progeny and recipients within the same pedigree structure are shown. (b) Geographical distribution of *SOC1*$^{\mathrm{Nip}}$-harboring cultivars across Japanese regions from the 1960s to 2010s. Each color represents a region from north to south in Japan. Areas delineated within each region correspond to individual prefectures.
